# Serotonin and dopamine modulate aging in response to food perception and availability

**DOI:** 10.1101/2021.03.23.436516

**Authors:** Hillary A. Miller, Shijiao Huang, Megan L. Schaller, Elizabeth S. Dean, Angela M. Tuckowski, Allyson S. Munneke, Safa Beydoun, Scott D. Pletcher, Scott F. Leiser

**Affiliations:** Cellular and Molecular Biology Program, University of Michigan, Ann Arbor, MI 48109, USA; Molecular & Integrative Physiology Department, University of Michigan, Ann Arbor, MI 48109; Department of Internal Medicine, University of Michigan, Ann Arbor, MI 48109, USA

**Keywords:** *C. elegans*, *D. melanogaster*, mammalian cells, lifespan, aging, mianserin, thioridazine, serotonin, dopamine, cell-nonautonomous, flavin containing monooxygenase, fmo-2, dietary restriction, nervous system

## Abstract

An organism’s ability to perceive and respond to changes in its environment is crucial for its health and survival. Here we reveal how the most well-studied longevity intervention, dietary restriction (DR), acts in-part through a cell non-autonomous signaling pathway that is inhibited by the perception of attractive smells. Using an intestinal reporter for a key gene induced by DR but suppressed by attractive smells, we identify three compounds that block food perception in *C. elegans*, thereby increasing longevity as DR mimetics. These compounds clearly implicate serotonin and dopamine in limiting lifespan in response to food perception. We further identify an enteric neuron in this pathway that signals through the serotonin receptor 5-HT1A/ser-4 and dopamine receptor DRD2/dop-3. Aspects of this pathway are conserved in D. melanogaster and mammalian cells. Thus, blocking food perception through antagonism of serotonin or dopamine receptors is a plausible approach to mimic the benefits of dietary restriction.

## Main

Rapid advances in aging research have identified several conserved signaling pathways that influence aging in organisms across taxa^1^. Recent work shows that many of these “longevity pathways” act through cell non-autonomous signaling mechanisms^2, 3^. These pathways utilize sensory cells—frequently neurons—to signal to peripheral tissues and promote survival during the presence of external stress. Importantly, this neuronal activation of stress response pathways, through either genetic modification or exposure to environmental stress, is often sufficient to improve health and longevity. Despite mounting evidence that neuronal signaling can influence multiple longevity pathways, less is known about which specific cells and molecules propagate these signals.

Biogenic amines are among the most well-studied and conserved neuronal signaling molecules^4,5^. Specifically, serotonin and dopamine play well-defined roles in behavior and physiology. However, their role in aging is less well understood. Several recent studies implicate serotonin, but not dopamine, as an important signal in multiple *C. elegans* longevity pathways including the response to heat shock and hypoxia^6, 7^. Dopaminergic signaling is associated with physical activity in humans and loss of this signaling decreases lifespan in mice^8^ and blocks lifespan extension in nematodes^9^. Serotonin and dopamine levels both decrease with age across species^10, 11^, consistent with these signaling pathways promoting healthy aging. Despite rigorous study and clinical use of drugs that modify serotonin and dopamine signaling, our understanding of their complex actions and potential interaction is far from complete.

Dietary restriction (DR) is the most well-studied and consistent intervention known to improve health and longevity in organisms ranging from single-celled yeast to primates^12^. DR leads to improved cell survival and stress resistance, complex intracellular signaling events, metabolic changes, and increased activity in multiple organisms. Nematode flavin-containing monooxygenase-2 (*fmo-2*) is necessary and sufficient to increase health and longevity downstream of DR. FMOs are highly conserved proteins that are also induced in multiple mammalian models with increased lifespan^13, 14^. Having previously identified a role for *fmo-2* in aging, we wondered whether DR cell non-autonomously regulates *fmo-2* induction and whether perception of food through biogenic amines could be involved in the subsequent signaling pathway.

## Results

### Attractant food perception represses *fmo-2* to limit longevity

We developed an integrated single-copy *mCherry* reporter driven by the *fmo-2* promoter to measure *fmo-2* induction. The reporter is primarily expressed in the intestine and responds to stimuli previously reported to induce *fmo-2*, including DR. As an intestinal protein^15^, we expected that *fmo-2* would likely be induced cell autonomously by the change in nutrient intake under DR. To test this hypothesis, we asked whether the perception of food smell by worms in the absence of eating can abrogate the induction of *fmo-2*. Using a “sandwich plate” assay as described in **Figure 1A**, we were surprised to find a significant reduction in *fmo-2* induction when worms could smell but not eat food (**Figure 1B-C**). This reduction is consistent with a model in which increased *fmo-2* mediates the increase in longevity from DR, as food smell completely abrogates this lifespan extension (**Figure 1D**, lifespan statistics in Table S1)^16^. We also find that active bacterial metabolism is required to abrogate *fmo-2* induction, as the “smell” of bacteria metabolically killed with 0.5% paraformaldehyde does not prevent DR from inducing *fmo-2* expression (Figure S1A-B). Since intestinal cells are not known to perceive external environmental cues such as smell, these results suggest that *fmo-2* expression is suppressed when food is present through cell non-autonomous signaling.

**Figure 1.**
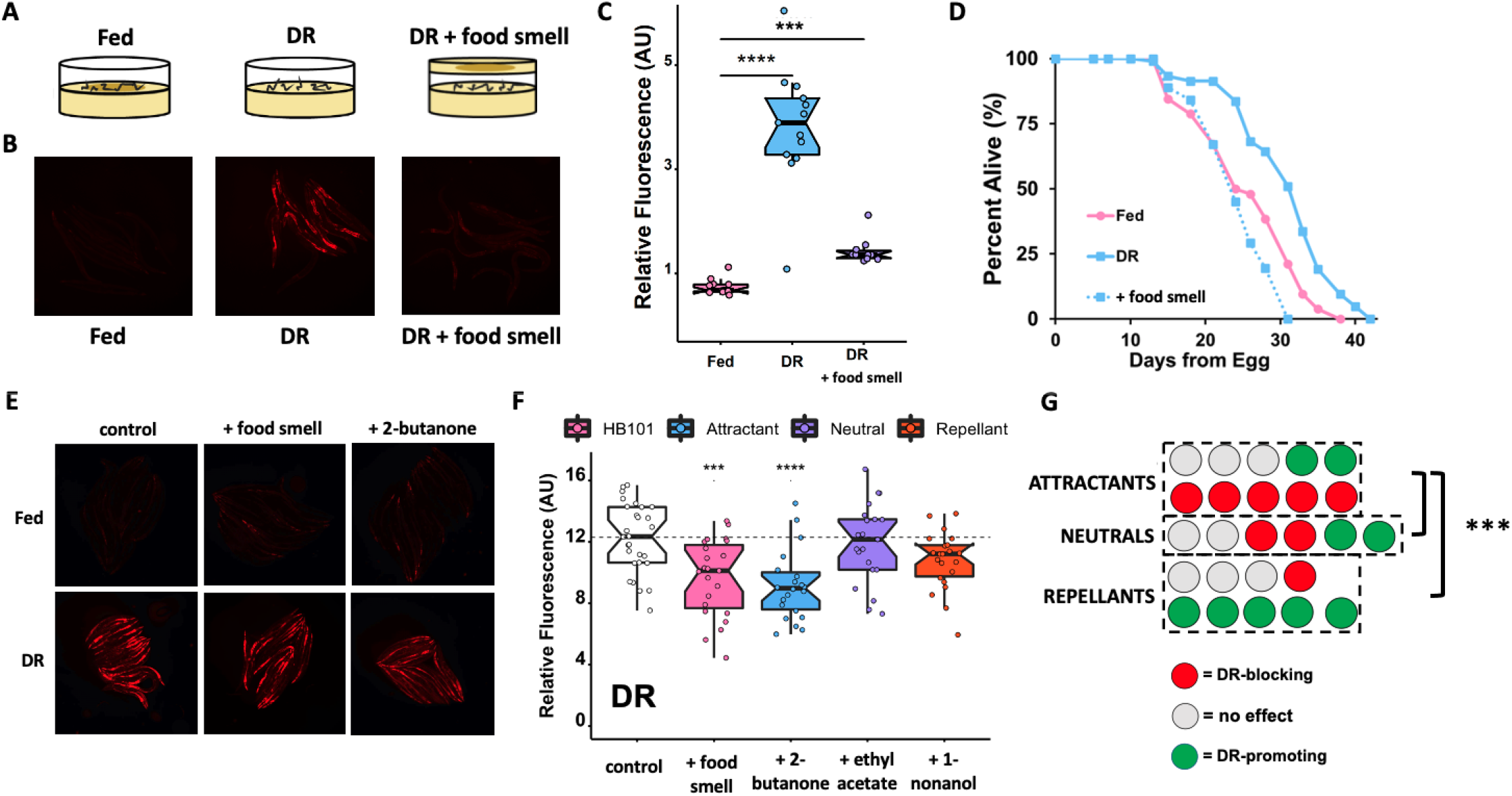
Attractive food smell blocks dietary restriction-mediated *fmo-2* induction and longevity. Diagram of “smell plates” (**A**). Images (**B**) and quantification (**C**) of individual *fmo-2p::mCherry* worms on fed (pink), DR (blue) and DR + food smell (OP50) (purple). Survival curves (**D**) of N2 (WT) animals fed (pink) or DR (blue) under normal conditions (solid lines) or subjected to the smell of bacteria (dotted lines). Images (**E**) and quantification (**F**) of individual *fmo-2p::mCherry* worms on DR plates exposed to food smell (HB101) (pink) or attractive (2-butanone in blue), neutral (ethyl acetate in purple), or repellant (1-nonanol in orange) odorants. (**G**) Summary of the effects of 26 odorants on *fmo-2* induction during DR. *** denotes P<.001, **** denotes P<.0001 when compared to fed (Tukey’s HSD) ### denotes P < 0.001 when compared to neutrals (ANOVA).

We next wondered what types of odorant compounds worms sense in this pathway. Bacteria are known to secrete hundreds of volatile compounds that are classified in three categories based on how they promote chemotaxis: attractants, repellants, and neutral compounds^17–19^. We tested whether exposure to any volatile compound secreted from bacteria is sufficient to block the lifespan-promoting effects of DR or whether compounds identified as attractants and repellants oppositely regulate *fmo-2* induction. Using compounds derived from studies of the *E. coli* strain HB101 in a range of concentrations (Table S2), we find that attractants are more likely to suppress DR-mediated induction of *fmo-2* (**Figure 1E-F**) whereas neutral and repellant compounds can induce *fmo-2* under fed conditions (Figure S1C-H). We also find that many compounds suppress *fmo-2* expression, consistent with the hypothesis that this pathway is not acting through a single receptor **(Figure 1G**, all results in Figure S2A-Z). These results support a model in which perception of attractive smells secreted by *E. coli* abrogates the induction of the pro-longevity gene *fmo-2*. This is consistent with these smells preventing the lifespan-promoting effects of DR, possibly through a neural response to external stimuli that leads to physiological changes in peripheral tissues.

### Serotonin and dopamine antagonists induce *fmo-2* to mimic DR longevity

Biogenic amines can regulate pro-longevity pathways and are involved in behavioral changes in response to food^6, 7, 20–22^. We next asked whether neurotransmitters are involved in the *fmo-*2-mediated food perception pathway. Using a targeted approach focusing on neurotransmitters and their antagonists, we tested for compounds sufficient to prevent the abrogation of *fmo-2* induction in the presence of food smell (Figure S3A-D). The biogenic amine neurotransmitter antagonists mianserin (for serotonin) and thioridazine and trifluoperazine (for dopamine) consistently and significantly restore *fmo-2* induction to DR levels in the presence of food smell (**Figure 2A-C**, Figure S3E-F). Mianserin is a tetracycline serotonin antagonist that is thought to competitively bind to specific serotonergic G protein-coupled receptors (GPCRs)^23^ while thioridazine and trifluoperazine’s mechanism of action involves blocking dopamine receptors^24^. Importantly, while each compound induces *fmo-2* to a different extent (**Figure 2D**, Figure S3Gand S3I), when combined with DR, no antagonist further induced *fmo-2*, suggesting they act in the same pathway (**Figure 2E**, Figure S3H-I). Diphenyleneiodonium chloride (DPI), an inhibitor of NADPH oxidase, acts as a positive control, and further induces *fmo-2* when combined with DR (**Figure 2E**). Because thioridazine and trifluoperazine act through similar mechanisms and the effects of thioridazine were more consistent in our studies, we focused further experiments on dopamine antagonism through thioridazine. Together, these results support antagonism of serotonin or dopamine as partial mimetics of DR in their induction of *fmo-2*.

**Figure 2.**
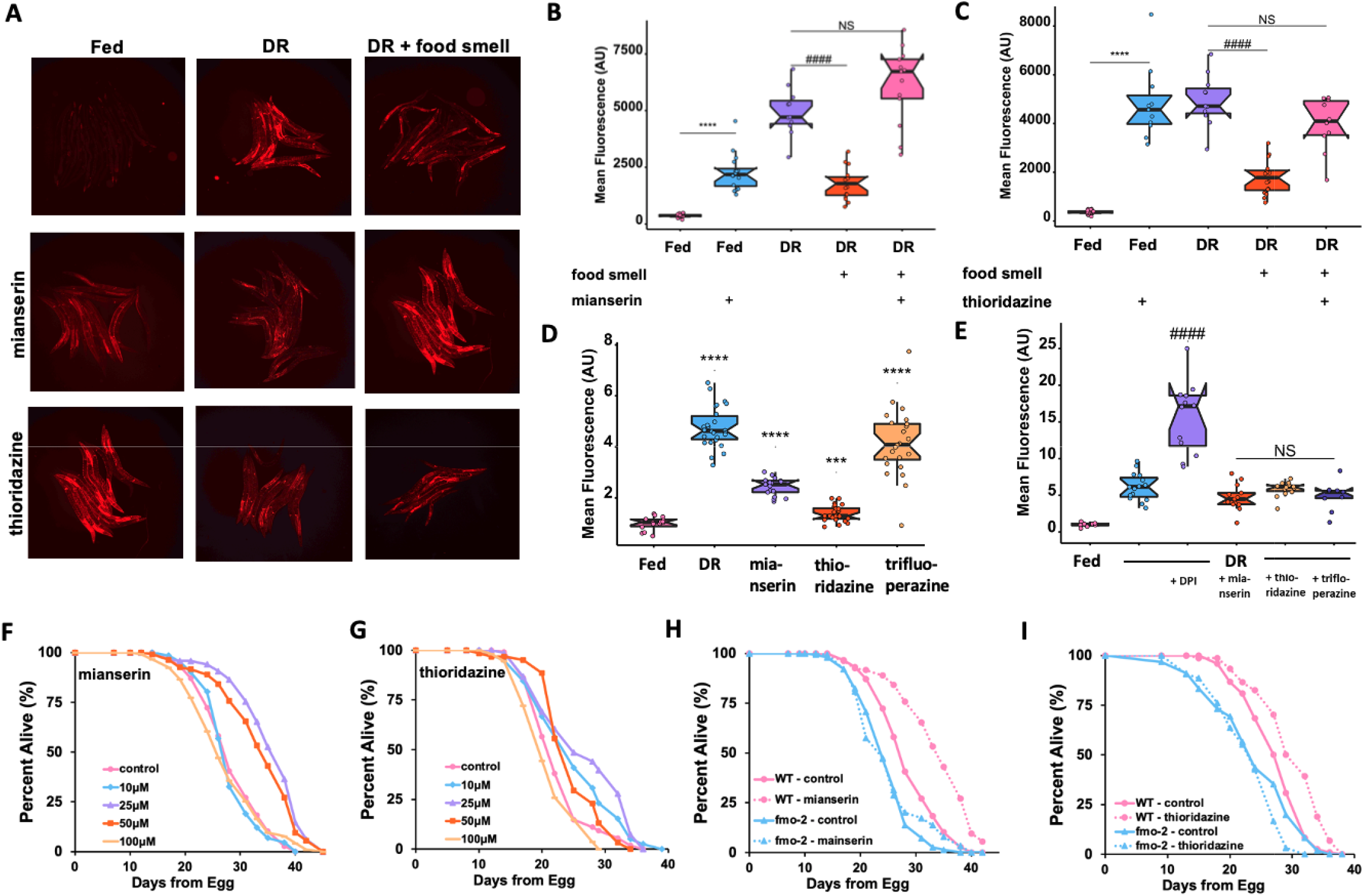
Serotonin and dopamine antagonists induce *fmo-2* and extend lifespan. Images (**A**) and quantification of *fmo-2p::mCherry* exposed 100μM of mianserin (**B**) or thioridazine (**C**) (blue) in combination with DR (orange) **and** food smell (pink) compared to DR alone (purple). Quantification (**D**) of *fmo-2p::mCherry* exposed to water (pink), DR (blue), 100μM mianserin (purple), thioridazine (orange), or trifluoperazine (yellow). Quantification (**E**) of *fmo-2p::mCherry* exposed to water (pink) or DR (blue) in combination with 500μM DPI (purple), 100μM mianserin (orange), 100**μM** thioridazine (yellow), or 100μM trifluoperazine (dark purple). Survival curves (**F**) of N2 (WT) animals treated w**ith** 0μM (water; pink), 10μM (blue), 25μM (purple), 50μM (orange), or 100μM (yellow) mianserin. Survival curves (**G**) of WT animals treated with 0μM (water; pink), 10μM (blue), 25μM (purple), 50μM (orange), or 100μM (yellow) thior**ida**zine. Survival curves (**H**) of WT animals (pink) and *fmo-2* KO animals (blue) on water (solid lines) or 50μM mianserin (dotted lines). Survival curves (**I**) of WT animals (pink) and *fmo-2* KO animals (blue) on water (solid lines) or 25μM thioridazine (dotted lines). *** denotes P<.001, **** denotes P<.0001 when compared to fed (Tukey’s HSD). #### denotes P<.0001 when compared to DR (Tukey’s HSD).

To validate that the induction of *fmo-2* through biogenic amine antagonism is beneficial for longevity, we next asked whether these compounds extend lifespan. We find that both mianserin and thioridazine extend lifespan on agar plates in a dose-dependent manner (**Figure 2F-G**)^25^. Since we identified mianserin and thioridazine through their induction of *fmo-2*, and previously found that *fmo-2* is necessary for DR-mediated lifespan extension, we next asked whether *fmo-2* was necessary for the beneficial longevity effects of mianserin or thioridazine. Our results show that the *fmo-2* loss of function completely blocks the lifespan effect of mianserin (**Figure 2H**) and thioridazine (**Figure 2I**). Importantly, we also see that mianserin treatment combined with DR does not further extend lifespan (Figure S3J). These results are consistent with these compounds mimicking some aspects of DR-signaling, recapitulating part of the DR lifespan extension effect. Collectively, this supports a model where DR induces *fmo-2* because of decreased biogenic amine signaling and establishes neuromodulators as a useful tool to decipher where in the signaling pathway a cell, signal, or receptor plays a role in DR-mediated longevity.

### DR signaling acts through a pair of enteric neurons

Our initial results establish that antagonizing serotonin and dopamine signaling leads to induction of the longevity promoting *fmo-2* gene and rescue of the negative effects of food smell. Based on this, we hypothesized that the relative lack of food smell during DR leads to increased longevity through induction of intestinal *fmo-2*. Using this framework, we next sought to better understand how the sensing of bacteria (or lack thereof) is communicated to intestinal cells during DR. Our results, knocking down the synaptic vesicle exocytosis gene *unc-13*, support short-range neurotransmitters as necessary for *fmo-2* induction (Figure S4A-B).

In *C. elegans*, perception of the external environment is largely regulated by a specialized organ known as the amphid. Since a previous report using a solid-liquid DR approach suggested a pathway originating in the ASI amphid neurons, we first asked whether these cells are required to modulate *fmo-2* activity during DR^26^. We find that not only are the ASI neurons (as measured by *daf-3* and *daf-7* RNAi) dispensable for food perception-mediated reduction in *fmo-2* expression (Figure S4C-D), but proper formation of the amphid (*daf-6*) is also not required (Figure S4E-F). This result is consistent with a non-canonical sensory neuron playing a role in food perception-mediated *fmo-2* suppression.

To better map this pathway, we next asked whether the biogenic amine serotonin is involved in the DR-mediated longevity pathway, and if so, where. We tested whether knocking out serotonin signaling would mimic the effects of DR. We subjected animals lacking *tph-1*, the rate-limiting enzyme necessary to produce serotonin, to DR and mianserin. *tph-1* animals are long-lived compared to wild-type^27^ and not further extended by our DR protocol (**Figure 3A**) or mianserin treatment (Figure S5A). These data are supported by the abatement of *fmo-2* induction on DR (**Figure 3B-C**) and mianserin (Figure S5B-C) when animals are subjected to *tph-1*(*RNAi*). As post-mitotic animals, *C. elegans* have a finite number of neurons with discrete connectivity and functions. Three neuronal pairs normally express *tph-1^28^*. The hermaphrodite specific motor neurons (HSN) are located along the ventral tail and regulate egg-laying^29^ whereas two head neuron pairs, the amphid neurons with dual sensory endings (ADF) and the neurosecretory motor (NSM) neurons, are involved in modifying behavioral states^20, 30, 31^. To investigate the potential role of these neuron pairs, we utilized *tph-1* cell-specific knockout and rescue strains and found that *tph-1* expression in NSM, but not the ADF, neurons (Figure S5D-E) is necessary (Figure S5F) and sufficient (**Figure 3D**) to promote DR-mediated longevity. These results implicate the NSM neurons as two of the primary neurons involved in reversing the effects of DR under food smell.

**Figure 3.**
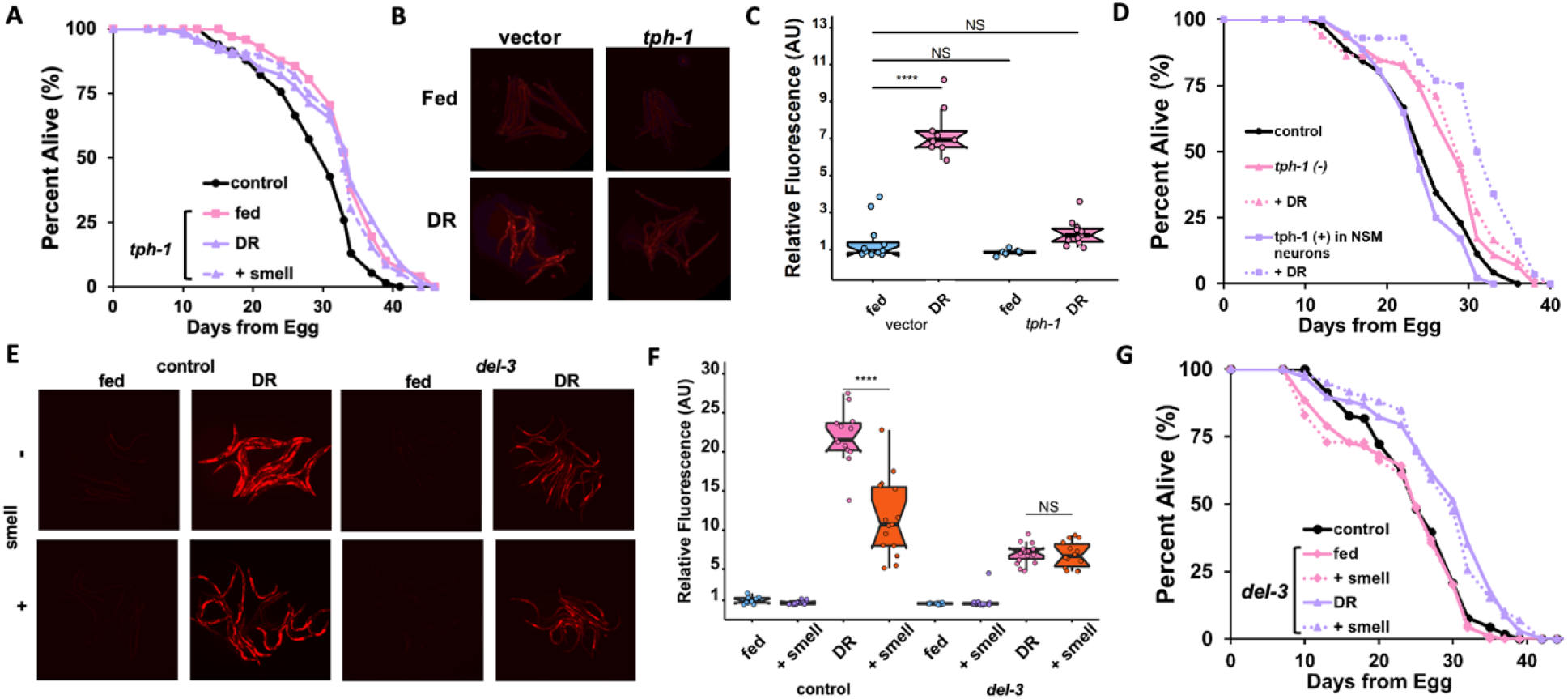
Food signals emanate from the NSM neurons. Survival curves (**A**) of WT animals (black) and *tph-1* KO animals on fed (pink) and DR (purple) conditions exposed to food smell (dotted lines). Images (**B**) and quantification (**C**) of *fmo-2p::mCherry* exposed to *tph-1* RNAi on fed (blue) or DR (pink). Survival curves (**D**) comparing control (black), *tph-1* KO (pink), and *tph-1* NSM-specific rescue (purple) animals on fed (solid line) and DR (dotted lines). Images (**E**) and quantification (**F**) of *fmo-2p::mCherry* in a WT (control) and *del-3* background on fed (blue) and DR (pink) exposed to food smell (purple and orange, respectively). Survival curves of conditions comparing WT (black) to *del-3* (**G**) on fed (pink) an**d** DR (purple) conditions in combination with food smell (dotted lines). **** denotes P<.0001 when compared to vector RNAi fed (Tukey’s HSD).

Recent research posits that NSM neurons function similar to enteric neurons with neural projections that directly communicate with the pharynx through a pair of acid-sensing ion channels (ASICs), DEL-3 and DEL-7. Signaling through these channels informs the worm to slow locomotion upon contact with food^30^. These data led us to wonder whether the longevity effects of DR also require the ASICs channels to extend lifespan. We find that *del-7* mutants look phenotypically wild type in their induction of *fmo-2* and lifespan extension, in either DR or DR + food smell (Figure S6A-C). Interestingly, *del-3* mutant worms show abrogated induction of *fmo-2* under DR, and did not diminish *fmo-2* induction in response to the smell of food (**Figure 3E-F**). These *del-3* mutant animals still exhibit lifespan extension under DR, despite the decreased induction of *fmo-2*, which is not abrogated by the smell of food (**Figure 3G**).

Together, these data support a model whereby the enteric NSM neurons release serotonin in response to food perception and the lack of this release extends longevity. In addition, the ASIC DEL-3 plays a role in the NSM to both behaviorally^30^ and physiologically respond to food perception signals.

### Mianserin mimics DR by antagonizing the 5-HT1A receptor SER-4

Prior reports suggest that serotonin receptor orthologs *ser-1* and *ser-4* are necessary for the lifespan benefits of mianserin in *C. elegans*^32^, and we hypothesized that a subset of the serotonin receptor orthologs will also be necessary for mianserin and DR-mediated *fmo-2* induction. After two generations of RNAi treatment, *ser-1* and *ser-4* were the only two receptors that proved necessary for *fmo-2* induction on mianserin (**Figure 4A**, Figure S7A-C) whereas *ser-4* knockdown most robustly abrogated DR-mediated *fmo-2* induction (Figure S7D-E). Further, we see that *ser-4(RNAi)* slightly but significantly increases lifespan and prevents DR from extending lifespan (**Figure 4B**), supporting the hypothesis that mianserin acts as a DR mimetic by antagonizing serotonin signaling that occurs during feeding. Finally, to investigate whether this effect is mediated by neuronal signaling or intestinal SER-4 expression, we rescued *ser-4* knockout animals with tissue-specific promoters and found that only neuronal *unc-119p::ser-4* is sufficient to rescue full induction of *fmo-2* under DR (**Figure 4C-D**). This is consistent with serotonergic signaling within the nervous system, and not directly to the intestine, regulating the response to food and food smell.

**Figure 4.**
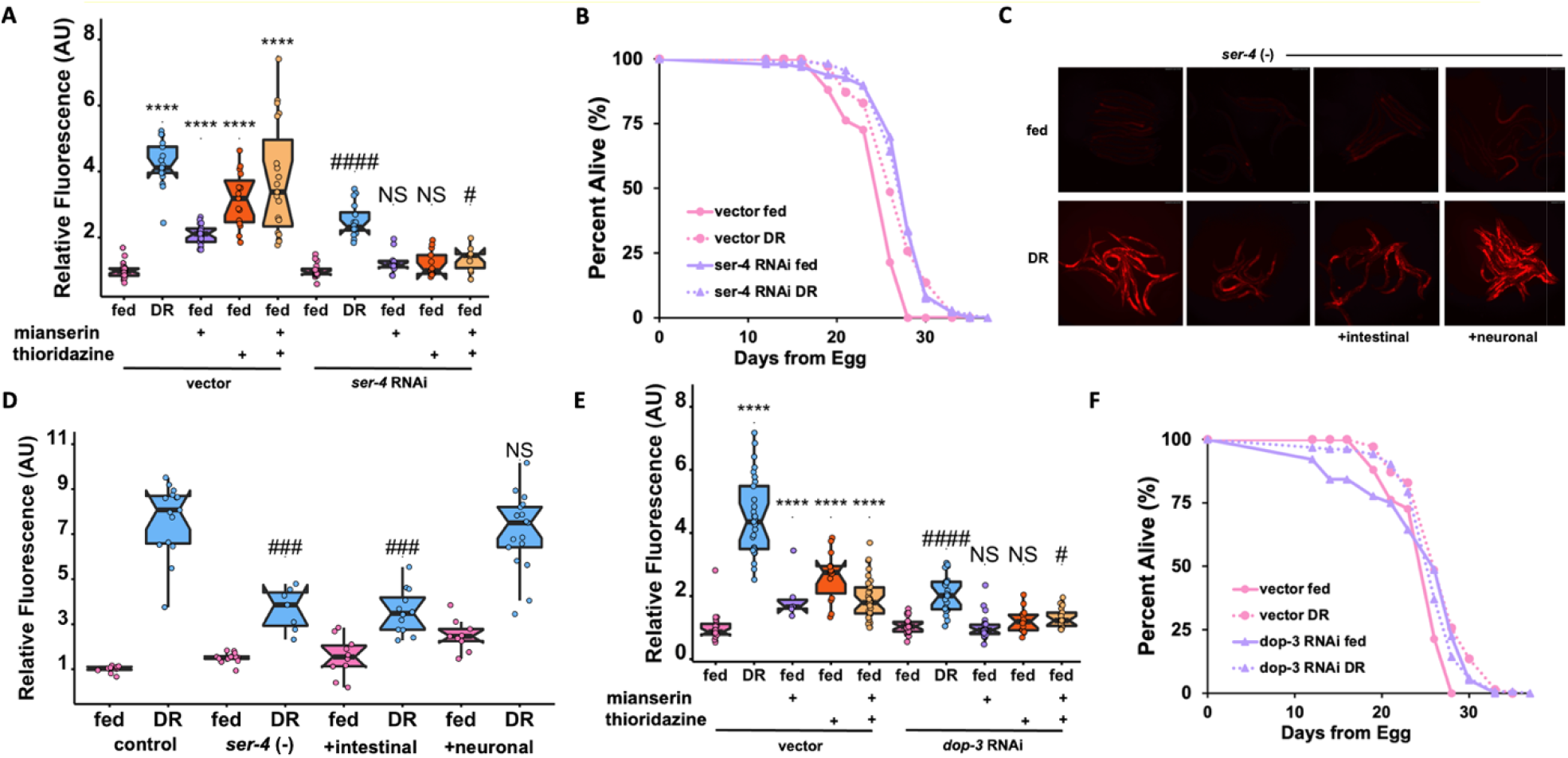
5-HT1A receptor *ser-4* and DRD2 receptor *dop-3* act downstream of food perception. Quantification (**A**) of individual *fmo-2p::mCherry* worms on fed (pink), and DR (blue) treated with 100μM mianserin (purple), 100 μM thioridazine (orange), or combined (orange) worms fed vector or *ser-4* RNAi. Survival curves (**B**) of WT animals on vector RNAi in pink and *ser-4* RNAi in purple on fed (solid lines) or DR (dotted lines) conditions. Images (**C**) and quantification (**D**) of *fmo-2p::mCherry* or *ser-4* with tissue-specific rescues added back on fed (pink) and DR (blue). Quantification (**E**) of individual *fmo-2p::mCherry* worms on fed (pink), and DR (blue) treated with 100μM mianserin (purple), 100 μM thioridazine (orange), or combined (orange) worms fed vector or *dop-3* RNAi. Survival curves (**F**) of WT animals on vector RNAi in pink and *dop-3* RNAi in purple on fed (solid lines) or DR (dotted lines) conditions. **** denotes P<.0001 when compared to vector RNAi fed (Tukey’s HSD). # denotes P<.05, #### denotes P<.0001 when compared to *ser-4/dop-3* RNAi fed (Tukey’s HSD).

### Thioridazine induces *fmo-2* and extends lifespan through Dopamine receptor DOP-3/DRD2

Thioridazine is a compound that antagonizes dopamine receptor D2 (DRD2) in mammals^33^, and induces *fmo-2* and mimics DR to increase longevity in nematodes (**Figure 2**). Based on its role in mammals, we tested whether nematode DRD2 is involved in DR and mianserin-related *fmo-2* induction and longevity. When the DRD2 ortholog *dop-3* is knocked down by RNAi, *fmo-2* induction is not affected in fed conditions but its induction by DR is diminished, while its induction by thioridazine is completely abrogated (**Figure 4E**, Figure S8A). This result is consistent with *dop-3* being required for dopaminergic induction of *fmo-2*. To demonstrate the epistasis of DOP-3 and SER-4 in the signaling pathway, we combined *ser-4* RNAi with mianserin and thioridazine treatment. The results show that *ser-4* depletion blocks *fmo-2* induction by thioridazine as well as suppresses *fmo-2* induction by mianserin, as expected (**Figure 4A**). Similarly, depletion of *dop-3* blocks both mianserin and thioridazine from inducing *fmo-2* (**Figure 4E**). These results support a model where both serotonin and dopamine signaling are epistatic to each other and are each required for full induction of *fmo-2* under DR. Interestingly, when *ser-4* or *dop-3* receptors are completely absent, via null mutation, the mutant animals show dysregulation of *fmo-2* induction, suggesting that the lack of biogenic amine signaling increases variability in responding to environmental changes (Figure S7F-G, S8B-D). To test whether DOP-3*/*DRD2 is necessary for lifespan extension by DR and mianserin, we depleted *dop-3* with RNAi under DR and found that *dop-3* depletion increases lifespan but is not further extended by DR (**Figure 4F**). Together, these results suggest that dopamine and serotonin signaling interactively induce *fmo-2* and extend lifespan under DR.

### Mianserin induces FMOs and promotes stress resistance in mammalian cells

Having identified serotonin and dopamine antagonism upstream of *fmo-2* induction under DR, we were curious whether these relationships might be conserved. In mammals, previous studies show interventions that increase longevity often both induced Fmo genes and increased stress resistance^13, 34^. Thus we tested whether mianserin and thioridazine are sufficient to induce mammalian Fmo genes and whether this induction could confer stress resistance, as a surrogate for longevity^35^. Our results, using human liver (HepG2) cells, show that while thioridazine did not lead to any changes (Figure S9A), perhaps due to lack of DRD2 receptor expression, mianserin treatment at 2μM increased protein levels of mammalian FMO2 (**Figure 5A-B**) and FMO1 (Figure S9B-C), while 0.1μM mianserin increased protein levels of FMO4 (Figure S9B and S9D). FMO3 and FMO5 protein levels are not changed upon mianserin treatment (Figure S9B and S9E-F). Since stress resistance is often correlated with increased lifespan both within and between species, and *fmo-2* increases stress resistance in *C. elegans*, we next examined whether mianserin also promotes stress resistance^35^. We treated cells with paraquat, an inducer of mitochondrial oxidative stress through increased production of the reactive oxygen species (ROS) superoxide, and find that 2 μM mianserin, the dose that showed maximal induction of FMOs, slightly but significantly improves the survival of HepG2 cells under an increasing dose of paraquat (Figure S9G). These data support serotonin antagonism as a conserved mechanism to induce Fmo expression and improve stress resistance.

**Figure 5.**
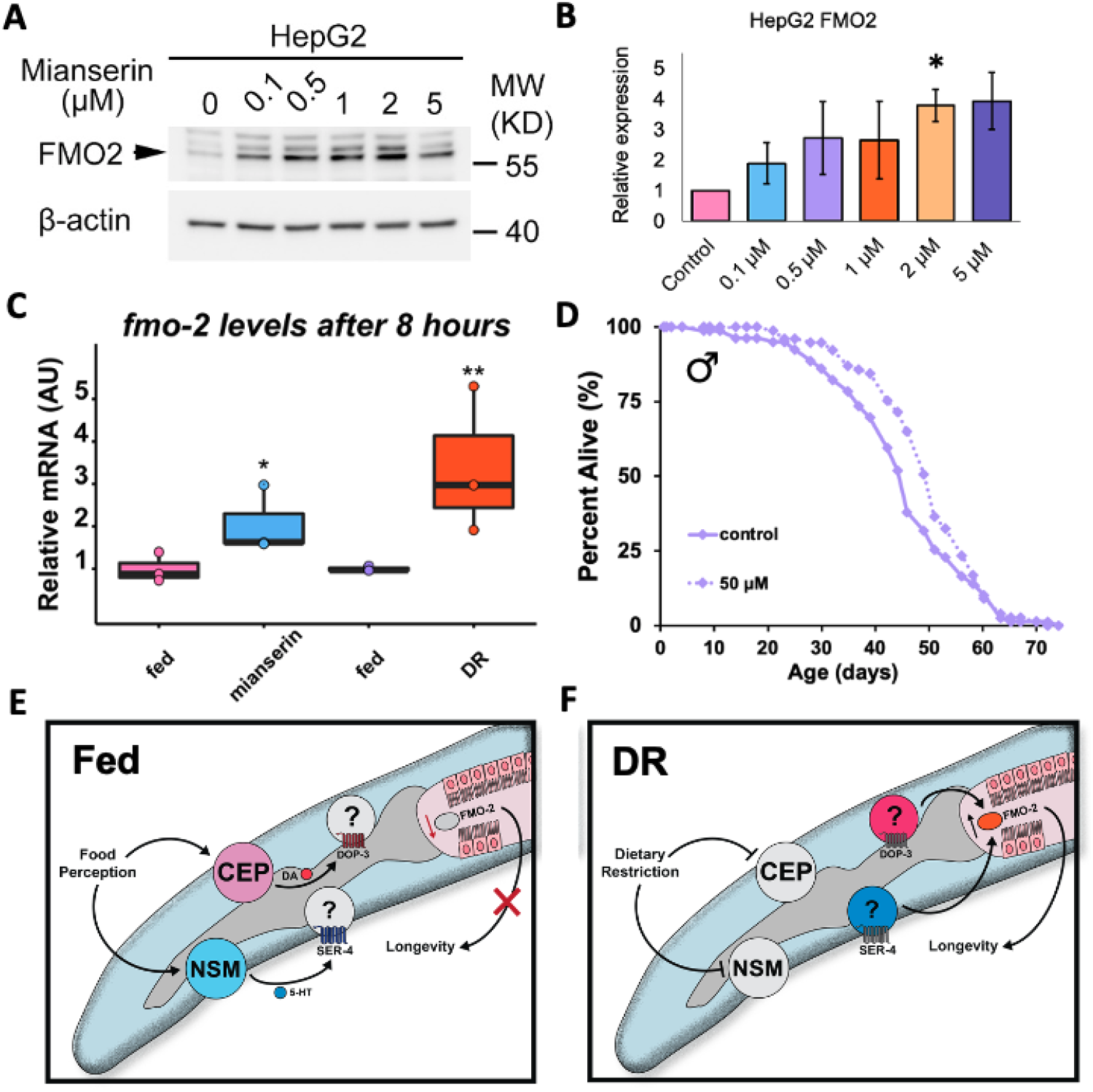
Serotonin antagonist mianserin induces FMO and improves health in Drosophila and mammalian cells. Western blot image (A) and quantification (B) of FMO1 in whole cell lysates from HepG2 cells treated with 0.1 μM, 0.5 μM, 1 μM, 2 μM, or 5 μM mianserin. Fmo-2 mRNA levels (C) after eight hours of 2mM mianserin (blue) or starvation (orange) compared to water controls (pink and purple, respectively). Survival curves of male flies treated with water (solid line) or 50μM (dotted line) mianserin (D). Panels E and F depict the “on/off” state worm’s toggle between when perceiving food.* denotes P<.05, ** denotes P<.01 when compared to control or fed (student’s t-test).

### Mianserin extends *D. melanogaster* lifespan similar to *C. elegans*

Since mianserin induces mammalian Fmos and promotes survival under paraquat stress, we tested whether it also affects lifespan in evolutionarily distant species. Similar to data in worms, recent data in the vinegar fly *D. melanogaster* show that altered serotonin signaling can change their ability to assess caloric quality and modulate lifespan^21^. As we found a narrow range of effective doses in worms (**Figure 2F**), we tested a slightly higher dose of mianserin in vinegar flies (2 mM) for its effect on Fmo2 induction. The resulting data show that both mianserin and fasting (DR) increase expression of fly *fmo-2* expression (**Figure 5C**), but not *fmo-1* (Figure S10A). We then asked whether mianserin could also extend lifespan in flies. Using several concentrations, we find a positive correlation between mianserin dosage and increased lifespan until reaching a detrimental level of serotonin antagonism (**Figure 5D**, Figure S10B-D). We also find a comparable dose response among male and female flies. We note that mianserin treatment does not significantly alter food consumption (Figure S10E-F), as measured by the Fly Liquid-Food Interaction Counter (FLIC) assay^36^. Together, these results are consistent with conserved induction of *fmos* by mianserin and DR, in addition to conserved lifespan extension.

## Discussion

Our experimental data in *C. elegans* support a model where the lack of an attractive (food) smell leads to a loss of serotonin release from the enteric NSM neurons and lack of serotonin binding to the SER-4/5-HT1A receptor. This in turn or in combination with other cues leads to a reduction in dopamine signaling to downstream DOP-3/DRD2 receptors. It is notable that both SER-4 and DOP-3 receptors are known to dampen adenylyl cyclase activity when bound, thus the lack of signal will increase the probability of excitement of the cell expressing these receptors. We hypothesize worms toggle their serotonin and dopamine neural activity “on” or “off” depending on the presence or absence of food, respectively (**Figure 5E-F**). Based on our ability to rescue DR benefits when food is perceived, we hypothesize that the perception of food during DR prevents the benefits of DR, rather than shortening lifespan through an independent pathway (Figure S10G). Critically, these data highlight that understanding how the nervous system evaluates and appropriately integrates large amounts of external stimuli, like the availability of food, allows us to target the decision-making processes to mimic pro-longevity pathways.

It is intriguing that dopamine and serotonin signaling interactively induce *fmo-2* and extend lifespan in a common pathway induced by dietary restriction. In nematodes, slowing locomotion in the presence of food is thought to be distinctly regulated by pharyngeal mechanosensation leading to dopamine release while dwelling behavior is potentiated by serotonin^37^. Significant scientific effort has identified much of the specific circuitry these neurotransmitters use to promote changes in chemotaxis and egg-laying^20, 30, 38–40^. The results suggest worms can interpret and implement a diverse set of responses to their changing environment. In mammals, SER-4/5-HT1A receptor activation increases dopamine release throughout the brain^41, 42^. Similarly, recent work shows release of serotonin and dopamine in the human brain influence non-reward-based aspects of cognition and behavior like decision making^43^. These findings support a conserved link between these two neurotransmitters in regulating complex phenotypes like aging.

It is also intriguing that one of these drugs, mianserin, successfully induces Fmo genes in both mammals and flies, and leads to increased stress resistance and lifespan, respectively. Since mianserin treatment extends fly lifespan we suspect it acts through a similar mechanism, serotonin antagonism, to mimic DR. This hypothesis is bolstered by *fmo-2* induction under acute mianserin exposure and fasting, analogous to what we see in *C. elegans*. It is not known whether FMOs or 5-HT1A receptors are necessary for mianserin or DR-mediated longevity in flies, but 5-HT2A receptors are necessary for proper food valuation^21^ suggesting that altering serotonin signaling may prove fruitful in future studies. In cells, the induction of Fmos by mianserin must be direct, suggesting that either serotonergic signaling is more direct in mammalian systems, or more likely, there are other nuances in this signaling in mammals we do not yet understand. Mammals and *C. elegans* share a single common ancestral Fmo^15^ and mammalian Fmos share similar homology to *C. elegans fmo-2*, with Fmo5 having the highest % identity. It is notable that 5-HT1A expression is detected in hepatocytes (The Human Protein Atlas), supporting a similar mechanism in these cells and suggesting that FMOs can be induced by serotonin antagonism both directly and indirectly. It will be interesting to investigate whether mianserin is beneficial for health and longevity in mammals. To achieve this goal, it is imperative that we understand the causative changes of pro-longevity drugs, such as atypical serotonin antagonists that are known to have pleiotropic effects in humans. In addition to providing the potential for long-term health benefits, this knowledge will benefit our understanding of serotonin and dopamine signaling networks that affect numerous human processes and diseases outside of aging.

## Authorship Contributions

H.A.M., S.H., and S.F.L developed the conceptual framework and wrote the manuscript.

H.A.M., S.H., M.L.S., E.S.D., A.M.T., A.S.M., S.B., and S.F.L. contributed to data collection and analysis. H.A.M. and E.S.D. prepared the figures and tables. All authors reviewed and approved the manuscript.

## Notes

### Competing Interest Statement

The authors have declared no competing interest.

## References

1. López-Otín C, Blasco MA, Partridge L, Serrano M, Kroemer G. The hallmarks of aging. Cell. Jun 2013;153(6):1194–217. doi:10.1016/j.cell.2013.05.039

2. Medkour Y, Svistkova V, Titorenko VI. Cell-Nonautonomous Mechanisms Underlying Cellular and Organismal Aging. Int Rev Cell Mol Biol. 2016;321:259–97. doi:10.1016/bs.ircmb.2015.09.003

3. Miller HA, Dean ES, Pletcher SD, Leiser SF. Cell non-autonomous regulation of health and longevity. Elife. Dec 2020;9 doi:10.7554/eLife.62659

4. Berger M, Gray JA, Roth BL. The expanded biology of serotonin. Annu Rev Med. 2009;60:355–66. doi:10.1146/annurev.med.60.042307.110802

5. Arias-Carrión O, Stamelou M, Murillo-Rodríguez E, Menéndez-González M, Pöppel E. Dopaminergic reward system: a short integrative review. Int Arch Med. Oct 2010;3:24. doi:10.1186/1755-7682-3-24

6. Tatum MC, Ooi FK, Chikka MR, et al. Neuronal serotonin release triggers the heat shock response in C. elegans in the absence of temperature increase. Curr Biol. Jan 2015;25(2): 163–74. doi:10.1016/j.cub.2014.11.040

7. Leiser SF, Miller H, Rossner R, et al. Cell nonautonomous activation of flavin-containing monooxygenase promotes longevity and health span. Science. Nov 2015;doi:10.1126/science.aac9257

8. Grady DL, Thanos PK, Corrada MM, et al. DRD4 genotype predicts longevity in mouse and human. J Neurosci. Jan 2013;33(1):286–91. doi:10.1523/JNEUROSCI.3515-12.2013

9. Saharia K, Kumar R, Gupta K, Mishra S, Subramaniam JR. Reserpine requires the D2-type receptor,. J Biosci. Dec 2016;41(4):689–695. doi:10.1007/s12038-016-9652-7

10. Yin JA, Liu XJ, Yuan J, Jiang J, Cai SQ. Longevity manipulations differentially affect serotonin/dopamine level and behavioral deterioration in aging Caenorhabditis elegans. J Neurosci. Mar 2014;34(11):3947–58. doi:10.1523/JNEUROSCI.4013-13.2014

11. Peters R. Ageing and the brain. Postgrad Med J. Feb 2006;82(964):84–8. doi:10.1136/pgmj.2005.036665

12. Fontana L, Partridge L, Longo VD. Extending Healthy Life Span--From Yeast to Humans. Science. April 16, 2010 2010;328(5976):321–326. doi:10.1126/science.1172539

13. Steinbaugh MJ, Sun LY, Bartke A, Miller RA. Activation of genes involved in xenobiotic metabolism is a shared signature of mouse models with extended lifespan. Am J Physiol Endocrinol Metab. Aug 2012;303(4):E488–95. doi:10.1152/ajpendo.00110.2012

14. Swindell WR. Genes and gene expression modules associated with caloric restriction and aging in the laboratory mouse. BMC Genomics. 2009;10:585. doi:10.1186/1471-2164-10-585

15. Petalcorin MI, Joshua GW, Agapow PM, Dolphin CT. The fmo genes of Caenorhabditis elegans and C. briggsae: characterisation, gene expression and comparative genomic analysis. Gene. Feb 2005;346:83–96. doi:10.1016/j.gene.2004.09.021

16. Kaeberlein TL, Smith ED, Tsuchiya M, et al. Lifespan extension in Caenorhabditis elegans by complete removal of food. Aging Cell. Dec 2006;5(6):487–94.

17. Bargmann CI, Hartwieg E, Horvitz HR. Odorant-selective genes and neurons mediate olfaction in C. elegans. Cell. Aug 1993;74(3):515–27. doi:10.1016/0092-8674(93)80053-h

18. Worthy SE, Haynes L, Chambers M, et al. Identification of attractive odorants released by preferred bacterial food found in the natural habitats of C. elegans. PLoS One. 2018;13(7):e0201158. doi:10.1371/journal.pone.0201158

19. Nuttley WM, Atkinson-Leadbeater KP, Van Der Kooy D. Serotonin mediates food-odor associative learning in the nematode Caenorhabditiselegans. Proc Natl Acad Sci U S A. Sep 2002;99(19):12449–54. doi:10.1073/pnas.192101699

20. Flavell SW, Pokala N, Macosko EZ, Albrecht DR, Larsch J, Bargmann CI. Serotonin and the neuropeptide PDF initiate and extend opposing behavioral states in C. elegans. Cell. Aug 2013;154(5):1023–35. doi:10.1016/j.cell.2013.08.001

21. Ro J, Pak G, Malec PA, et al. Serotonin signaling mediates protein valuation and aging. Elife. Aug 2016;5doi:10.7554/eLife.16843

22. Linford NJ, Ro J, Chung BY, Pletcher SD. Gustatory and metabolic perception of nutrient stress in Drosophila. Proc Natl Acad Sci U S A. Feb 2015;112(8):2587–92. doi:10.1073/pnas.1401501112

23. Marshall RJ. The pharmacology of mianserin--an update. Br J Clin Pharmacol. 1983;15 Suppl 2:263S–268S. doi:10.1111/j.1365-2125.1983.tb05874.x

24. Shana M. Feinberg KF, Abdolreza Saadabadi. StatPearls. Thioridazine. Stat Pearls; 2020.

25. Petrascheck M, Ye X, Buck LB. An antidepressant that extends lifespan in adult Caenorhabditis elegans. Nature. Nov 2007;450(7169):553–6. doi:10.1038/nature05991

26. Bishop NA, Guarente L. Two neurons mediate diet-restriction-induced longevity in C. elegans. Nature. May 2007;447(7144):545–9. doi:10.1038/nature05904

27. Murakami H, Murakami S. Serotonin receptors antagonistically modulate Caenorhabditis elegans longevity. Aging Cell. Aug 2007;6(4):483–8. doi:10.1111/j.1474-9726.2007.00303.x

28. Sze JY, Victor M, Loer C, Shi Y, Ruvkun G. Food and metabolic signalling defects in a Caenorhabditis elegans serotonin-synthesis mutant. Nature. Feb 2000;403(6769):560–4. doi:10.1038/35000609

29. Trent C, Tsung N, Horvitz HR. Egg-laying defective mutants of the nematode Caeenorhabditis Elegans. Genetics. August 1, 1983 1983;104(4):619–647.

30. Rhoades JL, Nelson JC, Nwabudike I, et al. ASICs Mediate Food Responses in an Enteric Serotonergic Neuron that Controls Foraging Behaviors. Cell. 01 2019;176(1-2):85–97.e14. doi:10.1016/j.cell.2018.11.023

31. Churgin MA, McCloskey RJ, Peters E, Fang-Yen C. Antagonistic Serotonergic and Octopaminergic Neural Circuits Mediate Food-Dependent Locomotory Behavior in Caenorhabditis elegans. J Neurosci. Aug 2017;37(33):7811–7823. doi:10.1523/JNEUROSCI.2636-16.2017

32. Petrascheck M, Ye X, Buck LB. A high-throughput screen for chemicals that increase the lifespan of Caenorhabditis elegans. Ann N YAcadSci. Jul 2009;1170:698–701. doi:10.1111/j.1749-6632.2009.04377.x

33. StatPearls. 2020.

34. Murakami S, Salmon A, Miller RA. Multiplex stress resistance in cells from long-lived dwarf mice. FASEB J. Aug 2003;17(11):1565–6. doi:10.1096/fj.02-1092fje

35. Harper JM, Salmon AB, Leiser SF, Galecki AT, Miller RA. Skin-derived fibroblasts from long-lived species are resistant to some, but not all, lethal stresses and to the mitochondrial inhibitor rotenone. Aging Cell. Feb 2007;6(1):1–13. doi:ACE255 [pii] 10.1111/j.1474-9726.2006.00255.x

36. Ro J, Harvanek ZM, Pletcher SD. FLIC: high-throughput, continuous analysis of feeding behaviors in Drosophila. PLoS One. 2014;9(6):e101107. doi:10.1371/journal.pone.0101107

37. Sawin ER, Ranganathan R, Horvitz HR. C. elegans locomotory rate is modulated by the environment through a dopaminergic pathway and by experience through a serotonergic pathway. Neuron. Jun 2000;26(3):619–31. doi:10.1016/s0896-6273(00)81199-x

38. Cermak N, Yu SK, Clark R, Huang YC, Baskoylu SN, Flavell SW. Whole-organism behavioral profiling reveals a role for dopamine in state-dependent motor program coupling in. Elife. Jun 2020;9doi:10.7554/eLife.57093

39. Hills T, Brockie PJ, Maricq AV. Dopamine and glutamate control area-restricted search behavior in Caenorhabditis elegans. J Neurosci. Feb 2004;24(5):1217–25. doi:10.1523/JNEUROSCI.1569-03.2004

40. Lee KS, Iwanir S, Kopito RB, et al. Serotonin-dependent kinetics of feeding bursts underlie a graded response to food availability in C. elegans. Nat Commun. 02 2017;8:14221. doi:10.1038/ncomms14221

41. Li Z, Ichikawa J, Dai J, Meltzer HY. Aripiprazole, a novel antipsychotic drug, preferentially increases dopamine release in the prefrontal cortex and hippocampus in rat brain. Eur J Pharmacol. Jun 2004;493(1-3):75–83. doi:10.1016/j.ejphar.2004.04.028

42. Bantick RA, De Vries MH, Grasby PM. The effect of a 5-HT1A receptor agonist on striatal dopamine release. Synapse. Aug 2005;57(2):67–75. doi:10.1002/syn.20156

43. Bang D, Kishida KT, Lohrenz T, et al. Sub-second Dopamine and Serotonin Signaling in Human Striatum during Perceptual Decision-Making. Neuron. Oct 2020;doi:10.1016/j.neuron.2020.09.015

