## Supplementary material for "Serotonin and dopamine modulate aging in response to food perception and availability": Methods + supplemental figures: Supplemental_Miller_Huang_etal_2021.docx

**Strains and growth conditions**

Standard procedures for C. elegans strain maintenance^1^ were used where experiments were performed on animals fed Escherichia coli (OP50) from egg and maintained on solid nematode growth medium (NGM). Additionally, worms were exposed to the smell of OP50 or HB101. Supplementary Table 3 includes a list of the strains and RNAi conditions used in this study. All genotypes were confirmed using PCR.

***fmo-2p::mCherry* construct**

We PCR amplified *mCherry* from pHG8 and the *fmo-2* promoter from the worm gDNA under the *fmo-2* promoter and cloned them into pdonr221 and P4-P1r, respectively. From here, they were combined using Gateway LR cloning (Invitrogen) to create *fmo-2p*::*mCherry*::unc-54 3’UTR on PCFJ150 (pHAM001).

**SER-4 rescue constructs**

We purchased donor plasmid pPD117.01 from Addgene and used Gibson cloning (NEB) to swap out promoters driving cDNA of *ser-4*::SL2::GFP (on backbone) expression. We used the *unc-119* promoter (pHAM002) to target all neurons and the *vha-6* promoter (pHAM003) to target the intestine. All plasmids were verified via restriction digest and sanger sequencing. ApE files available upon request.

**Microinjection**

Single-copy integration of pHAM001 using the ttTi5605 (EG6699) Mos allele was performed as previously described^2^. Overexpression transgenic animals were generated by injecting PureLink (Invitrogen) miniprepped DNA clones (~50ng/µL) with fluorescent co-injection marker *myo-2p*::mNeonGreen (15 ng/µL) and junk DNA (up to 100 ng/µL) into gonads of day 1 gravid adult hermaphrodites. Standard protocols were followed to isolate and obtain stable over-expression mutants^3^. Because transgene expression can vary substantially, we typically characterized 2-4 independent transgenic lines per experiment.

**Lifespan measurements**

Lifespans were carried out as previously described with minor modifications^4^. Briefly, 20 - 30 gravid adult animals were placed on NGM plates for a timed egg-lay. After 12-16 hours, these animals were removed. Once their progeny reached late L4/early adult stage, animals were transferred to plates with 33 µL of 150 mM fluorodeoxyuridine (FUdR) and 100 µL of 50 mg/mL Ampicillin per 100 mL NGM to prevent the development of progeny and growth of bacteria. Roughly 75 worms were placed on each NGM + FUdR plate seeded with concentrated bacteria (10×). A minimum of two plates per strain per condition were used per replicate. Lifespan plates were transferred periodically during early adulthood to prevent starvation and avoid contamination. Animals were scored as dead and removed from the experiment when they did not move in response to prodding under a dissection microscope.

**RNAi knockdown**

The RNAi feeding bacteria were obtained from the Vidal RNAi library. All RNAi plasmids were sequenced to verify the correct target sequence. Animals were exposed to RNAi plates from egg on plates consisting of NGM supplemented with 1 mM β-D-isothiogalactopyranoside (IPTG) and 25 μg/ml carbenicillin. At late L4 stage of development the animals were transferred to plates containing freshly seeded RNAi bacteria plus FUdR.

**PFA treatment**

In order to metabolically kill OP50 in food smell assays, bacteria cultures were treated with 0.5% PFA. After 16 hours of shaking, 50 mL of the bacteria were aliquoted into 250 mL Erlenmeyer flasks. 32% PFA was added to the flasks to get the desired final PFA concentration (e.g., 390 µL of PFA was added to get the final concentration of 0.25% PFA). PFA-treated bacteria were shaken at 37^o^C for 1 hour and then transferred to 50 mL conical tubes, centrifuged and washed with LB five times to remove residual PFA before seeding.

**Drug treatments**

Recent reports show improved health outcomes and longevity in nematodes treated with mianserin^5^, but only in liquid culture^6^. As our studies are on agar plates, we modified previous protocols by adding mianserin, thioridazine or trifluoperazine before pouring NGM agar plates. Without proper dosing, these neurotransmitter antagonists can cause off-target effects like fleeing, especially when combined with DR. All subsequent *C. elegans* experiments were performed at 50 µM of mianserin and 25 µM of thioridazine unless otherwise noted. All drugs were purchased from Sigma-Aldrich and were initially dissolved in milliQ water at 2 mM (mianserin) or 100mM concentration (DRD2 antagonists), aliquoted, and stored at −20 °C.

**Dietary restriction (DR) lifespan treatments**

Lifespan DR assays were performed like other lifespans until day two of adulthood, when the worms were transferred to plates with 10^9 seeded lawns and transferred every other day four times. This form of DR is termed solid DR (sDR) (Greer et al 2009). For short-term DR assays, worms were starved for eight (real-time PCR) or 20 hours (slide microscopy). We added 100uL of 10mM palmitic acid (Sigma-Aldrich) dissolved in 100% EtOH to the outer rim of the plate to prevent fleeing.

**Attractant, repellant, and neutral smell treatments**

Fed and DR plates were prepared using NGM plates with palmitic acid. Odorants were chosen from previously published work isolating secreted compounds from the *E. coli* strain HB101^7, 8^. All concentrations of attractant, repellant, and neutral chemicals were dissolved in 100% ethanol (more details in Table S2). A small pad of NGM agar (2 mL) was poured on the lid of each plate and allowed to solidify before 100uL of each smell concentration was added to the agar pads. Plates were prepared the day prior to use to allow the ethanol solutions to dry. Young adult *fmo-2p::mCherry* worms were placed on fed and DR plates and exposed to each smell for 20 h before fluorescent microscopy images were taken.

**Slide microscopy**

All images in this study were acquired using LASx software and Leica scope with >15 worms/treatment at 6.3x magnification. Worms were paralyzed in 0.5M sodium azide (NaN3). Fluorescence mean comparisons were quantified in ImageJ using the polygon tool and saved as macros.

**Real-time PCR**

500 N2 worms per biological replicate were transferred at young adulthood, 2.5 days post-hatch, to FuDR plates either seeded with food, DR’ed, or poured with the addition of 50µM mianserin or thioridazine. Worms were harvested in 50uL of M9 and flash frozen in liquid N2 after eight hours of exposure. Samples were freeze-thawed three times in Trizol reagent (Invitrogen) and RNA was extracted following standard phenol-chloroform protocols from the manufacturer. Superscript reverse transcriptase II (Invitrogen) was used to synthesize cDNA. 600ngs of cDNA/sample were used with PowerUp SYBR Green Master Mix (Applied Biosystems) was used in the quantitation with primers:

fmo-2 FWD ACGAAACGAATGAGTCGTCAGT; REV AGAGCAGACAAGAACGCCAT

Canton-S flies were mated and reared on standard food for 2 weeks before separating the flies by sex onto SY10 food with 20 flies/vial. Flies were acclimated to the vials for 24 hours before being transferred to SY10 vials coated with 2mM mianserin or water (control) or vials containing 2% agar to mimic dietary restriction. After 8 hours on these treatments, flies were frozen at -80°C overnight. Fly heads and bodies were then separated by vortexing and dissection by forceps (all samples and materials were kept on dry ice throughout). Each treatment contained 3 biological replicates composed of 10 bodies each. Trizol Reagent (Invitrogen) was used in the RNA extraction, the MultiScribe Reverse Transcriptase kit (Applied Biosystems) was used to synthesize the cDNA, and the real-time PCR analysis used PowerUp SYBR Green Master Mix (Applied Biosystems) and a StepOne Plus Real-time PCR system (Applied Biosystems) primers:

fmo-1 FWD GCGATAGGATGGGCAAACTG; REV CCCGGAAGTGGAGCAAATTC

fmo-2 FWD CGCAACCAGAAGAAAGCACA; REV TGCTCCTGTACGTGTCCAAT

**Fly husbandry**

The laboratory stock Canton-S was used in the lifespan and molecular experiments. Flies were maintained on standard food and housed at 25°C and 60% relative humidity in a 12:12 hour light-dark cycle.

**Fly survival assays**

For lifespan measurements, flies were reared under controlled larval density and collected onto standard food within 24 hours of eclosion. Flies were mated for 2-3 days then sorted by sex under light CO_2_ onto vials containing standard food used in lifespan experiments (10% sucrose/10% yeast, or SY10), according to well-establish lifespan protocols^9^. Flies were transferred to fresh food every 2-3 days. At the beginning of the lifespan, mianserin was dissolved in water at a 1mM stock concentration and stored at -20°C. Weekly aliquots were prepared and diluted with water to yield the final concentrations of 20-80µM. 100µL of the drug solution (or water for the control) was added to the top of each vial and kept at room temperature to dry for approximately 2 hours before transferring the flies.

**Cell culture and stress resistance assay**

HepG2 cells were grown in Dulbecco’s modified Eagle’s medium (DMEM, GIBCO) supplemented with 10% fetal bovine serum,100 U/ml penicillin, and 100 μg/ml streptomycin. For stress resistance assay, cells were seeded to 96-well microplates with 40,000 cells per well for HepG2 cells. After 16 to 18 h overnight incubation in complete medium, the cells were incubated for 18 to 24 h in serum-free DMEM supplemented with 2% bovine serum albumin (BSA) as described previously^10^. For stress treatments, cells were exposed to cadmium for 6 h in 2% BSA supplemented DMEM, and then incubated in fresh 2% BSA supplemented DMEM without stressor for 18 h, followed by measurements of cell survival by Cell Proliferation Reagent WST-1 (Sigma 5015944001).

**Western blot analysis**

10 μg cell lysis samples were separated using SDS–PAGE and transferred to nitrocellulose filters, then blocked with 3% milk in TTBS (20mM Tris–HCl [pH 7.4], 500mM NaCl and 0.1% Tween 20) for 1 h, and incubated with primary antibody overnight at 4°C. After washing three times with TTBS buffer, membrane was incubated with horseradish peroxidase-conjugated secondary antibody (Cell Signaling Technology, diluted 1: 5,000 in 3% milk) for 1 h at room temperature and then washed with TTBS. The filter was developed for visualization by enhanced chemiluminescence (Thermo Scientific Pierce).

**Statistical analyses**

All box plots show individual data points while the box represents SEM (centered on the mean), and whiskers represent 10%/90%. Comparisons between more than two groups were done using ANOVA. For multiple comparisons, Tukey’s multiple comparison test was used, and p values are *p < 0.05, **p < 0.01, ***p < 0.001, and ****p < 0.0001. For lifespan assays groupwise and pairwise comparisons among survivorship curves were performed the statistical software R. P values were obtained using the log-rank analysis (select pairwise comparisons and group comparisons or interaction studies) as noted. Interaction P values were calculated using Cox regression when the survival data satisfied the assumption of proportional hazards. All statistics were run in R. Summary lifespan data and statistics are included in Table S1.

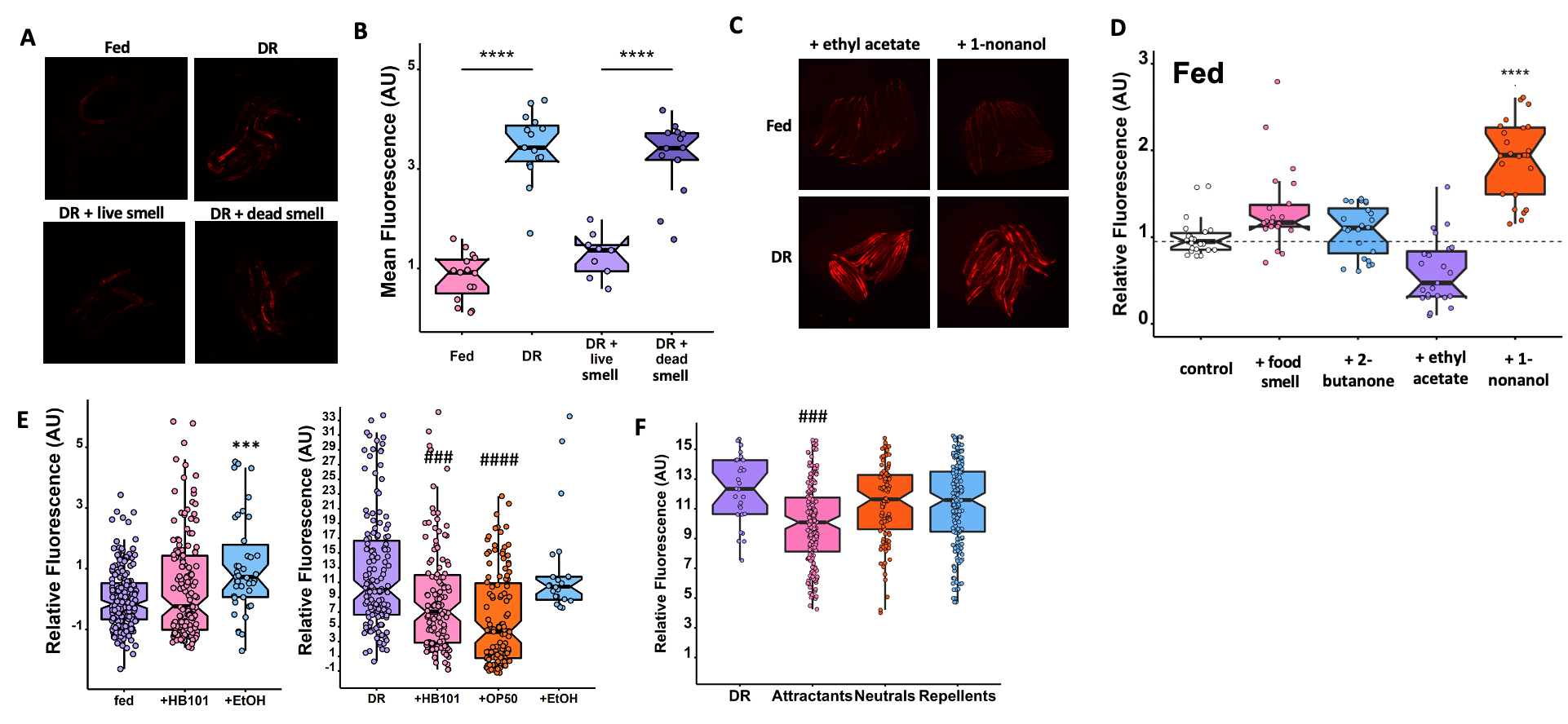

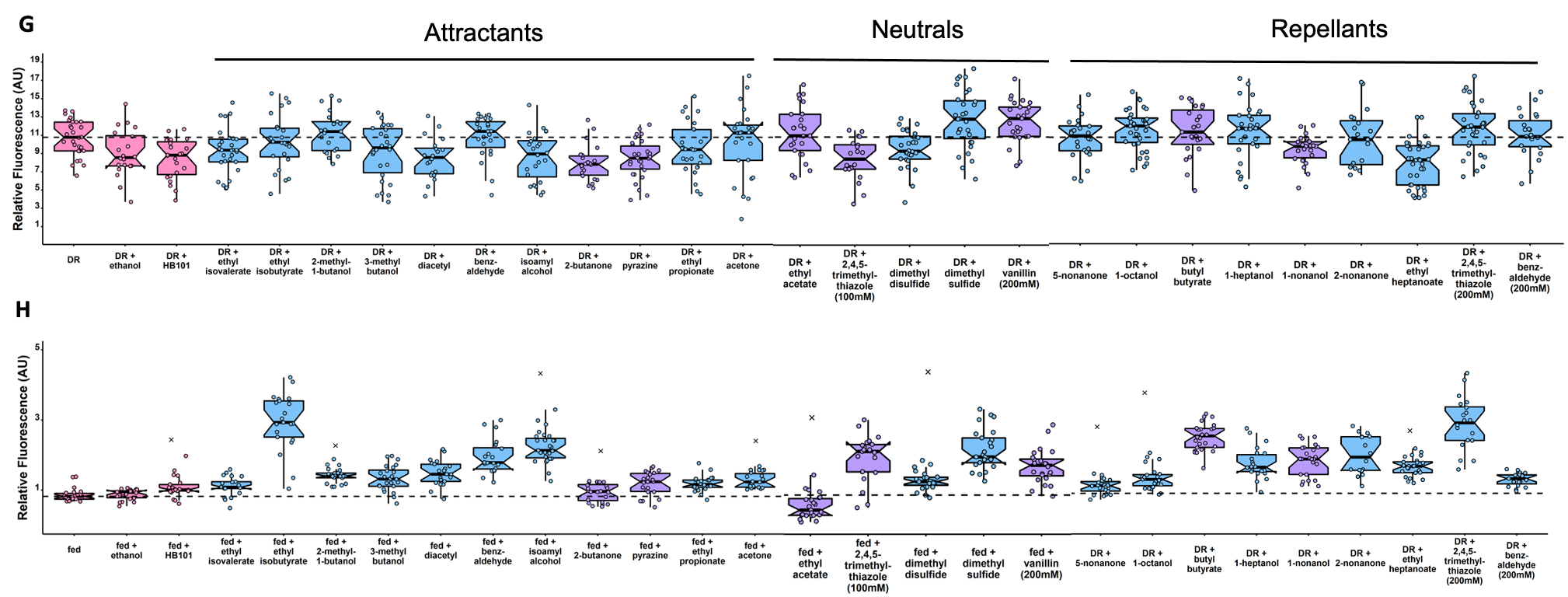

**Figure S1. Odorant effects on *fmo-2* expression.** Images (A) and quantification (B) of individual *fmo-2p::mCherry* worms on fed (pink), DR (blue), and smell with live OP50 (light purple) or PFA-killed OP50 (dark purple). Additional images (C) and quantification (D) of *individual fmo-2p::mCherry* worms on fed plates exposed to food smell (pink) or attractive (2-butanone in blue), neutral (ethyl acetate in purple), or repellant (1-nonanol in orange) odorants. Summary of controls HB101 (pink), OP50 (orange) and ethanol (blue) effects on fed and DR conditions across experiments (E). Summary of worms treated with attractant (pink), neutral (orange) or repellant (blue) compounds compared to DR (G). All odorants effects on DR (G) and fed (H) conditions. **** denotes P<.0001 when compared to fed (Tukey’s HSD). ### denotes P<.001 when compared to Neutrals or Repellants (Tukey’s HSD).

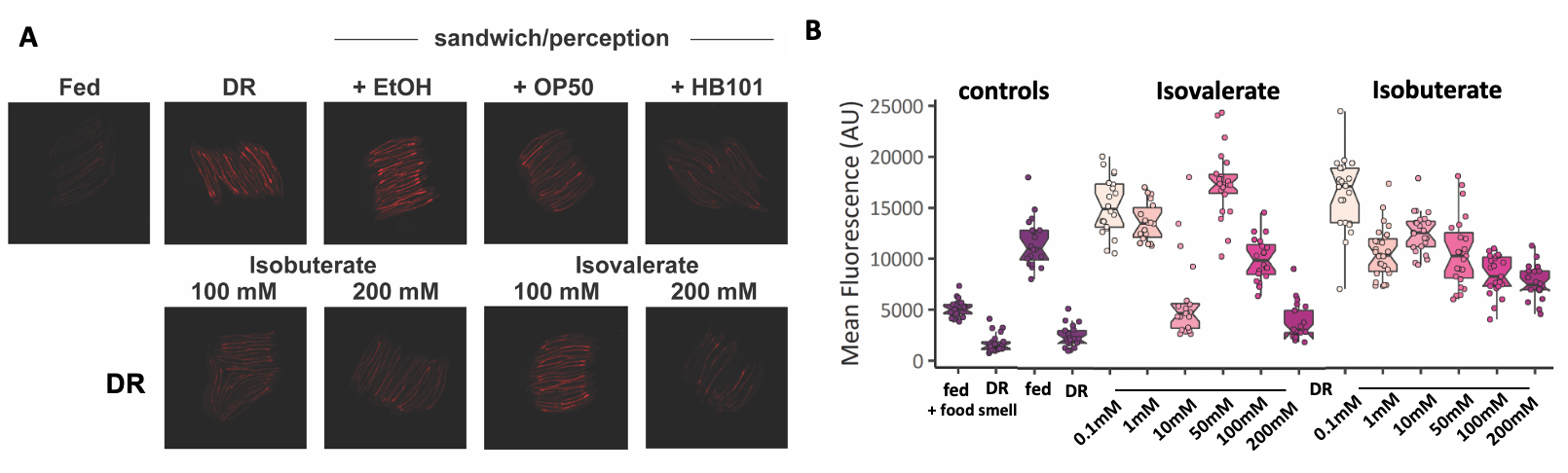

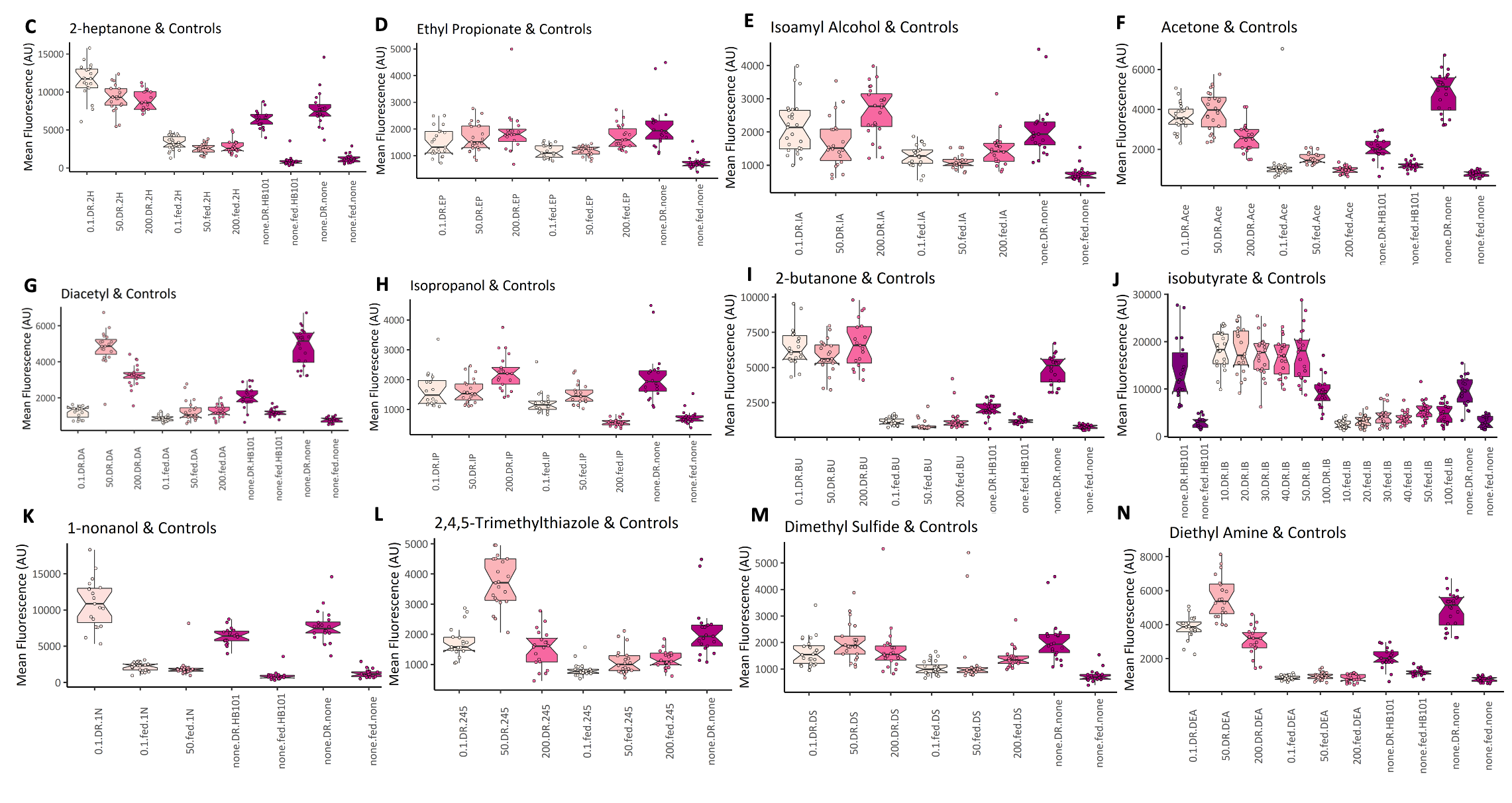

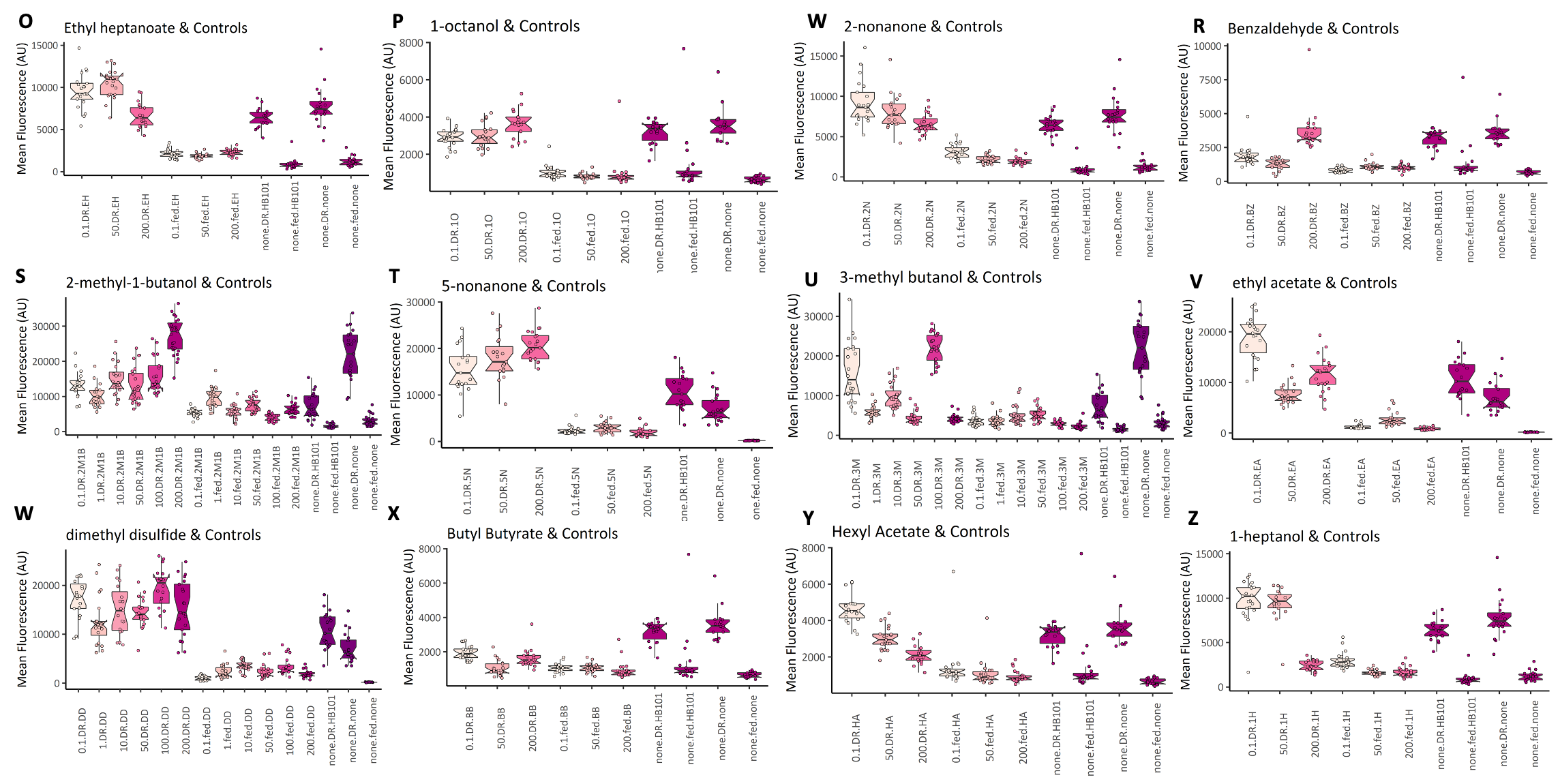

**Figure S2.** **Titration experiments of odorants tested**. Panels show representative images (A) and quantification of *fmo-2p::mCherry* under DR (B- Z). Dosing and preparation can be found in Table S2.

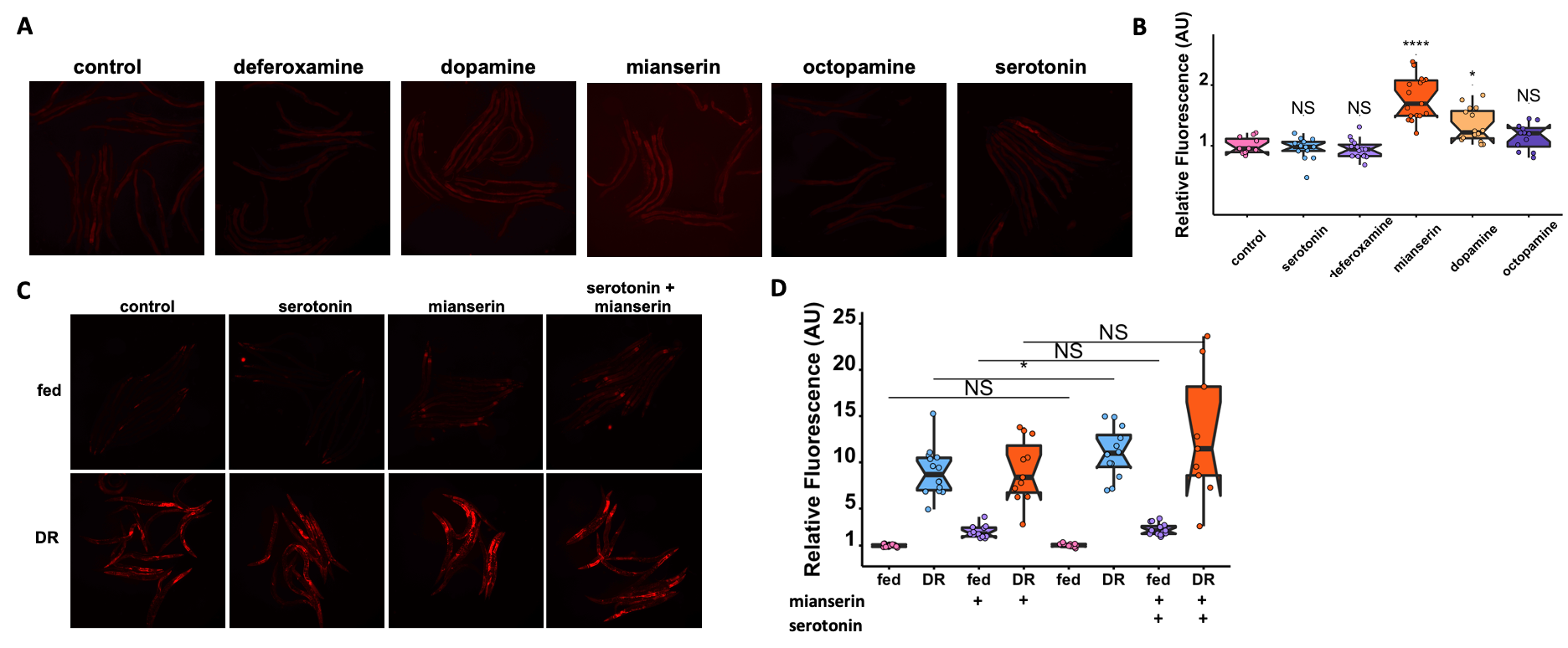

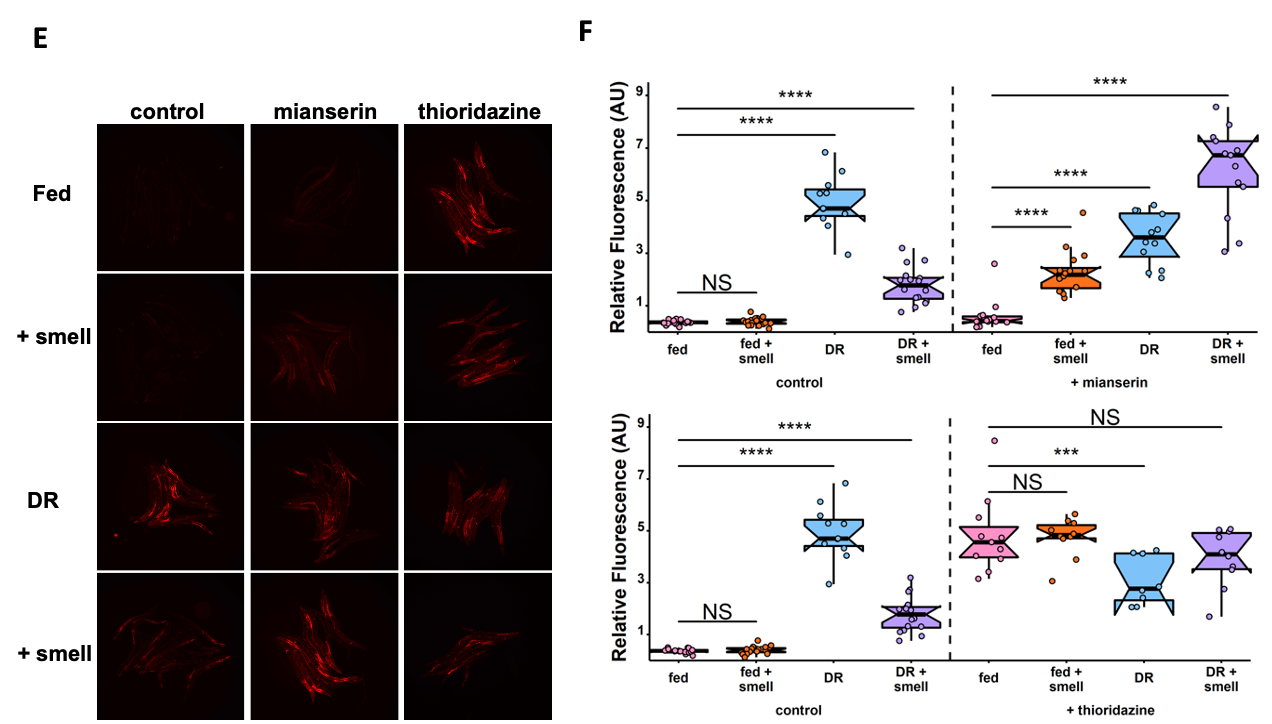

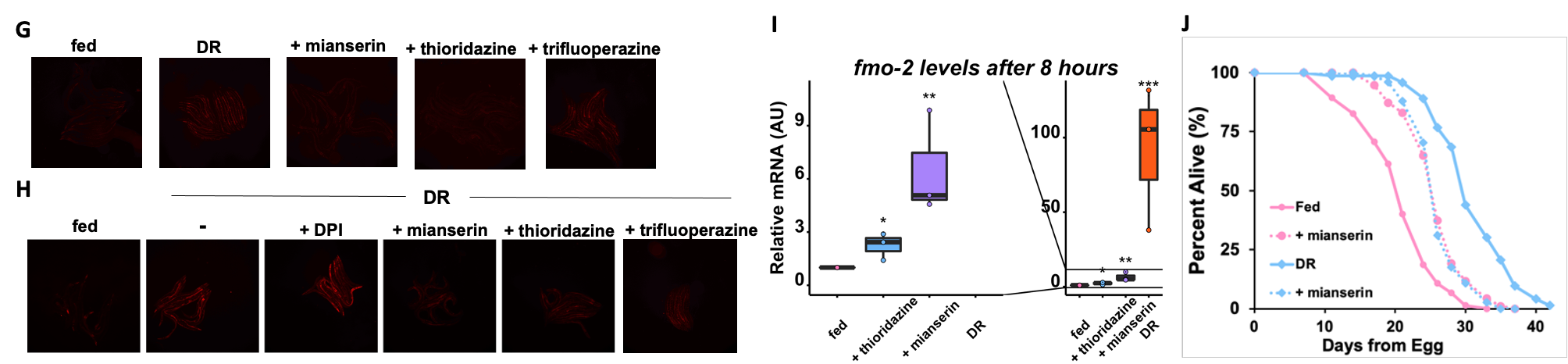

**Figure S3**. **Induction of *fmo-2* by neuromodulators.** Images (A) and quantification (B) of *fmo-2p*::*mCherry* worms exposed to water (pink), dopamine (blue), serotonin (purple), mianserin (orange), octopamine (yellow), or deferoxamine (dark purple). Images (C) and quantification (D) of *fmo-2p*::*mCherry* exposed to water (pink), DR (blue), mianserin (purple) or both (orange) in combination with serotonin. Additional control images (E) and quantification (F) from Fig 2A-C. Images (G) quantified in Figure 2d. Images (H) quantified in Figure 2E. qPCR results (I) for fmo-2 mRNA levels after 8 hours post DR (blue), mianserin (purple) or thioridazine (orange) treatment normalized to water control. Survival curves (J) of WT animals on fed conditions in pink and DR conditions in blue on water (solid lines) or 50µM mianserin (dotted lines). * denotes P<.05, ** denotes P<.01, **** denotes P<.0001 when compared to fed (Tukey’s HSD).

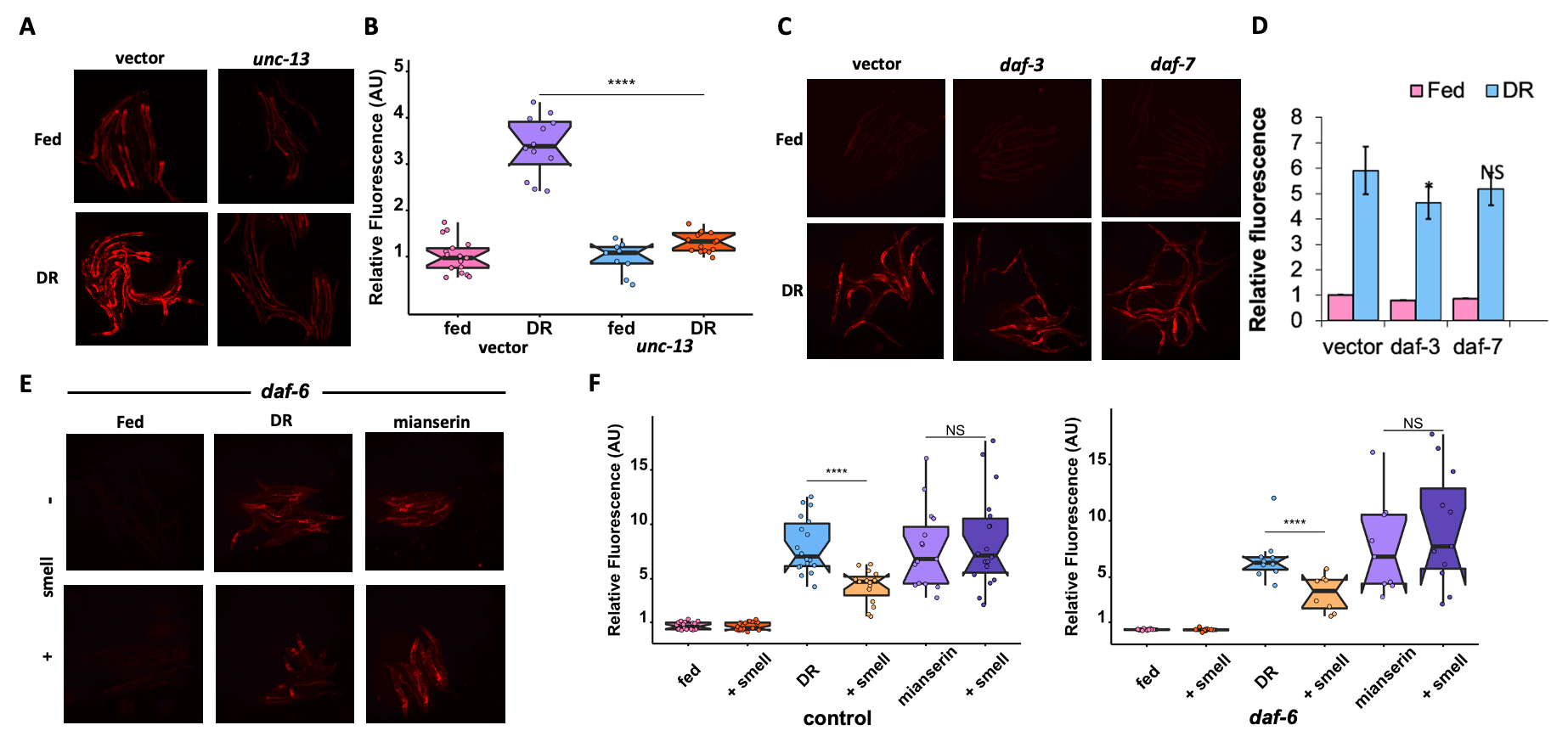

**Figure S4. Neuronal gene necessity for *fmo-2* induction under DR/food smell/biogenic amin antagonism.** Images (A) and quantification (B) of individual *fmo-2p::mCherry* worms on fed (pink, blue) and DR (purple, orange) fed vector or *unc-13* RNAi, respectively. Images (C) and quantification (D) of individual *fmo-2p::mCherry* worms on vector, *daf-3*, and *daf-7* RNAi on fed (pink) or DR (blue). Images (E) and quantification (F) of *fmo-2p*::*mCherry* in a *daf-7* KO background on fed, DR or exposed to mianserin (pink) and food smell (blue). * denotes P<.05, ** denotes P<.01, *** denotes P<.001, **** denotes P<.0001 when compared to DR (Tukey’s HSD).

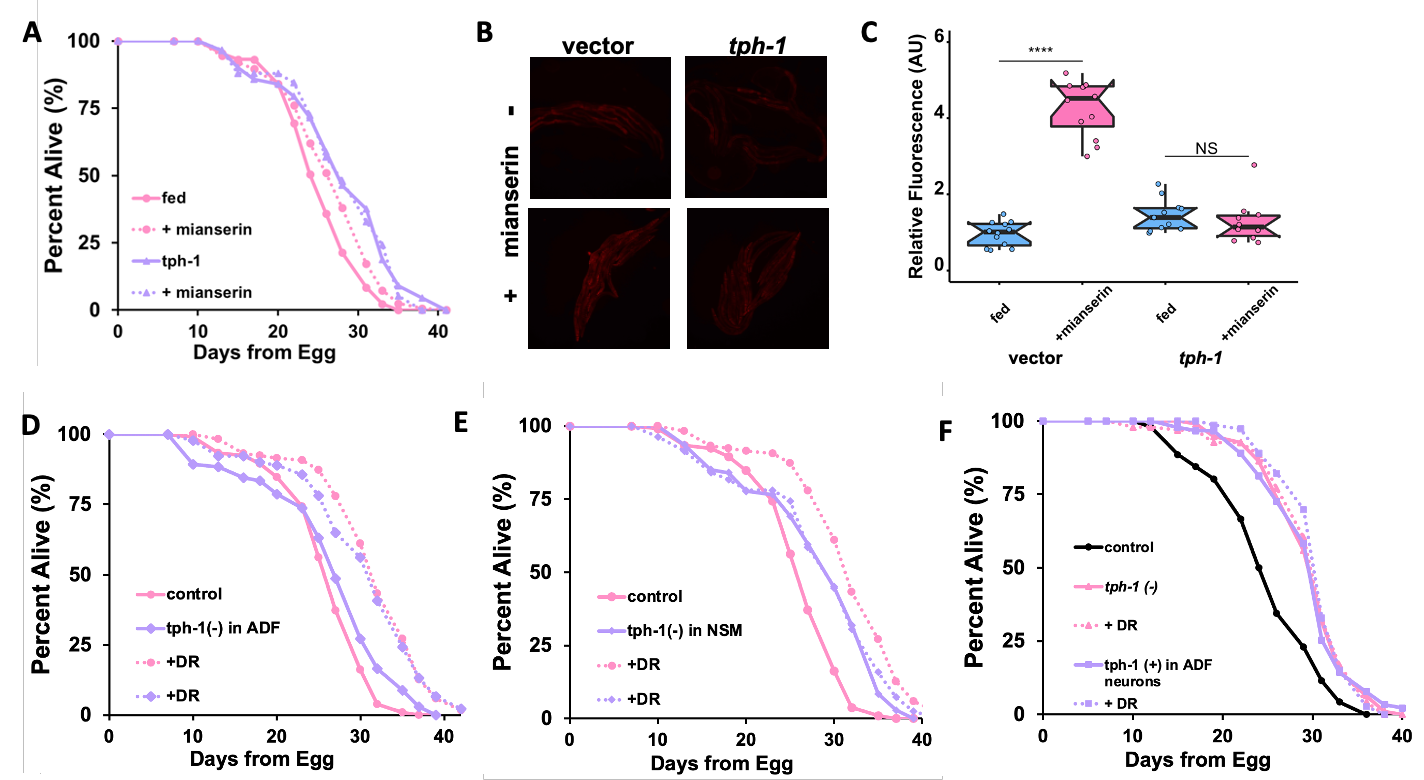

**Figure S5. Serotonin and serotonergic neuron-regulation of *fmo-2* induction and longevity.** Survival curves (A) of WT animals in pink and *tph-1* KO animals in purple on water (solid lines) or 50µM mianserin (dotted lines). Images (B) and quantification (C) of individual *fmo-2p::mCherry* worms on *tph-1* RNAi exposed to water (pink) or 50µM mianserin (blue) conditions. Survival curves comparing control (pink) and *tph-1* NSM-specific (D) or ADF-specific KO (E) (purple) animals on fed (solid line) and DR (dotted lines). Survival curves (F) comparing control (black), *tph-1* KO (pink), and *tph-1* ADF-specific rescue (purple) animals on fed (solid line) and DR (dotted lines). **** denotes P<.0001 compared to fed (Tukey’s HSD).

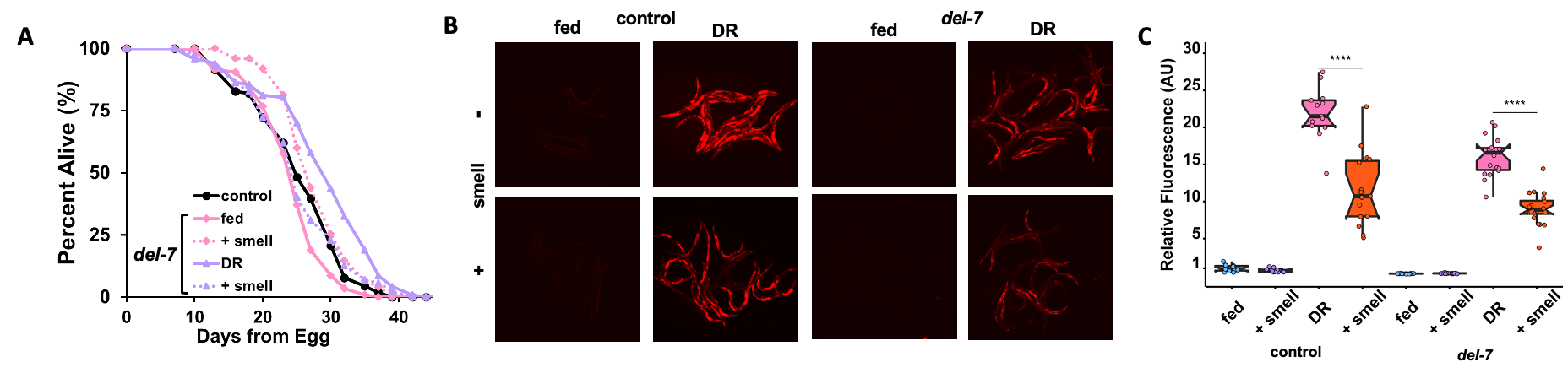
**Figure S6. ASICs channels modify responses to DR and food smell.** Survival curves of conditions comparing WT (black) to *del-7* (A) on fed (pink) and DR (purple) conditions in combination with food smell (dotted lines). Images (B) and quantification (C) of *fmo-2p*::*mCherry* in a WT (control) and *del-7* background on fed (blue) and DR (pink) exposed to food smell (purple and orange, respectively). **** denotes P<.0001 compared to DR (Tukey’s HSD).

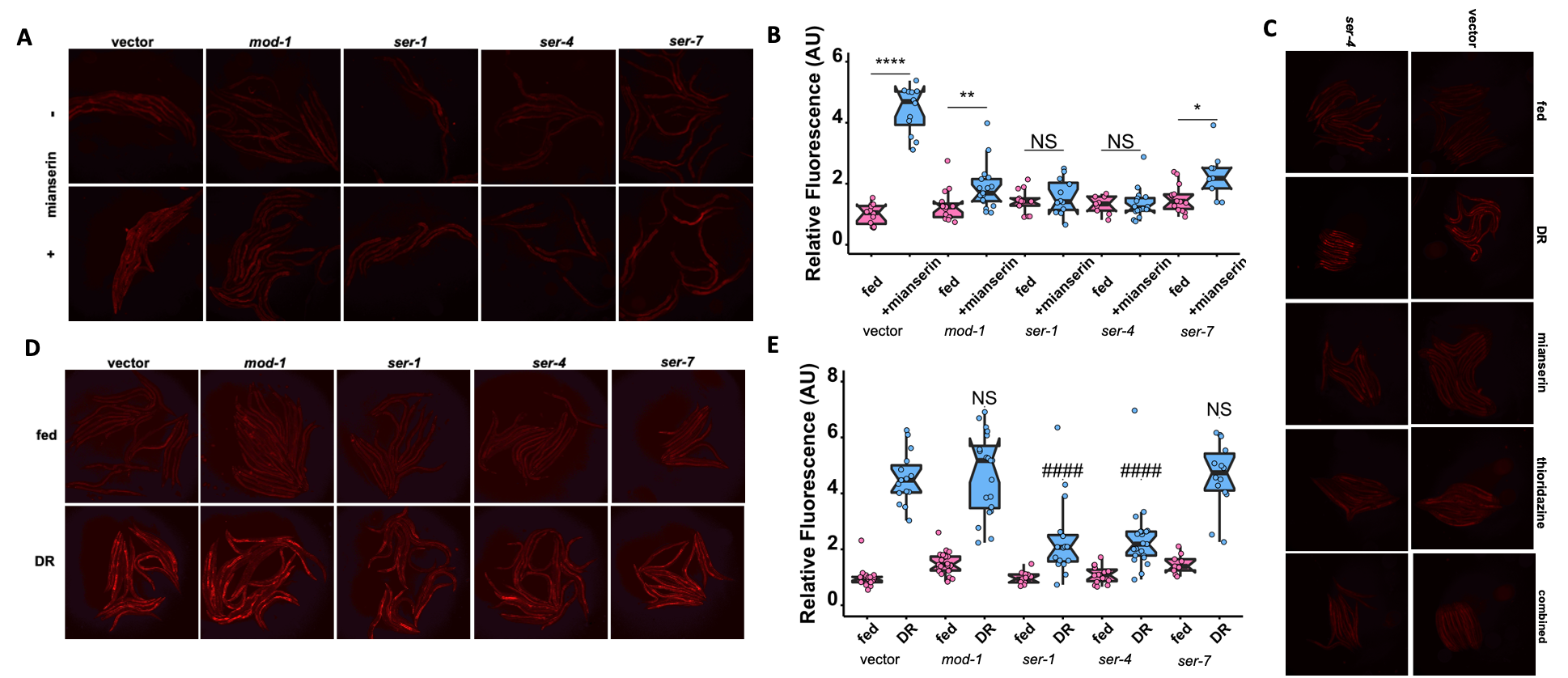

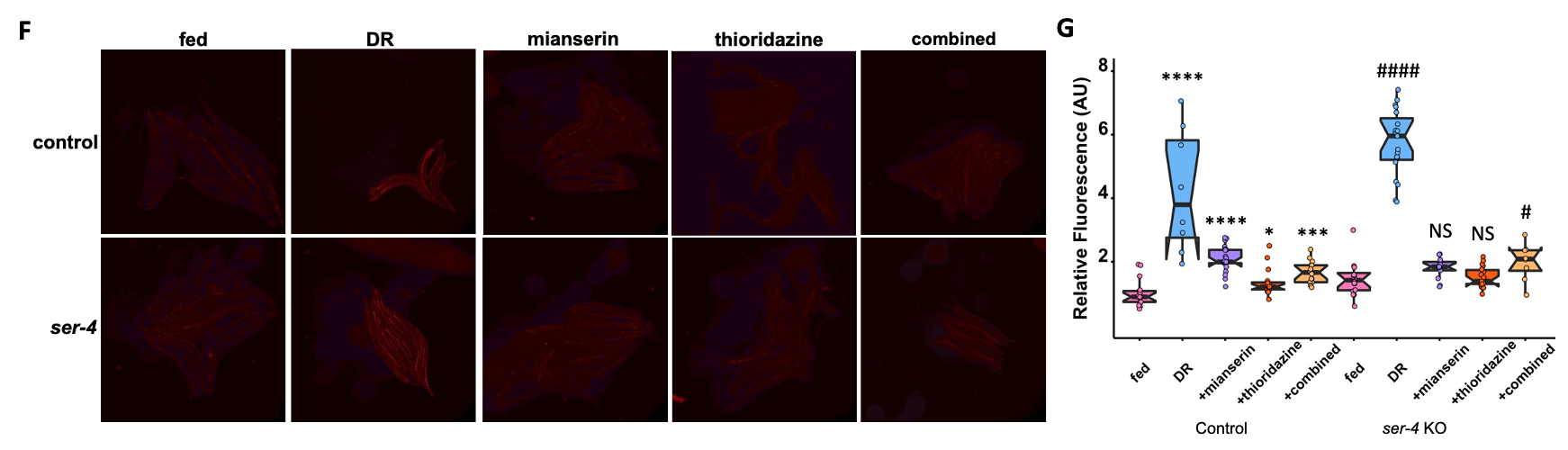

**Figure S7.** **The role of serotonergic receptor signaling in *fmo-2* induction by DR and DR mimetics.** Images (A) and quantification (B) of *fmo-2p*::*mCherry* grown on serotonin receptor RNAi exposed to water (pink) or 50µM mianserin (blue). Images (C) quantified in Figure 3E. Images (D) and quantification (E) of *fmo-2p*::*mCherry* grown on serotonin receptor RNAi exposed to fed (pink) or DR (blue). Images (F) and quantification (G) of WT *fmo-2p*::*mCherry* or *ser-4* KO on fed (pink), and DR (blue) treated with 100µM mianserin (purple), 100 µM thioridazine (orange), or combined (orange). * denotes P<.05, ** denotes P<.01, **** denotes P<.001 when compared to fed (Tukey’s HSD). ### denotes P<.001, #### denotes P< .0001 when compared to DR (Tukey’s HSD).

**Figure S8. The role of dopaminergic receptor signaling in *fmo-2* induction and lifespan extension by DR and DR mimetics.** Images (A) quantified in Figure 3g. Summary (B) of fed and DR conditions of WT and *ser-4* KO animals across multiple experiments. Images (C) and quantification (D) of WT *fmo-2p*::*mCherry* or *dop-3* KO on fed (pink), and DR (blue) treated with 100µM mianserin (purple), 100 µM thioridazine (orange), or combined (orange). * denotes P<.05, ** denotes P<.01, *** denotes P<.001 when compared to fed (Tukey’s HSD). ### denotes P<.001 when compared to DR (Tukey’s HSD).

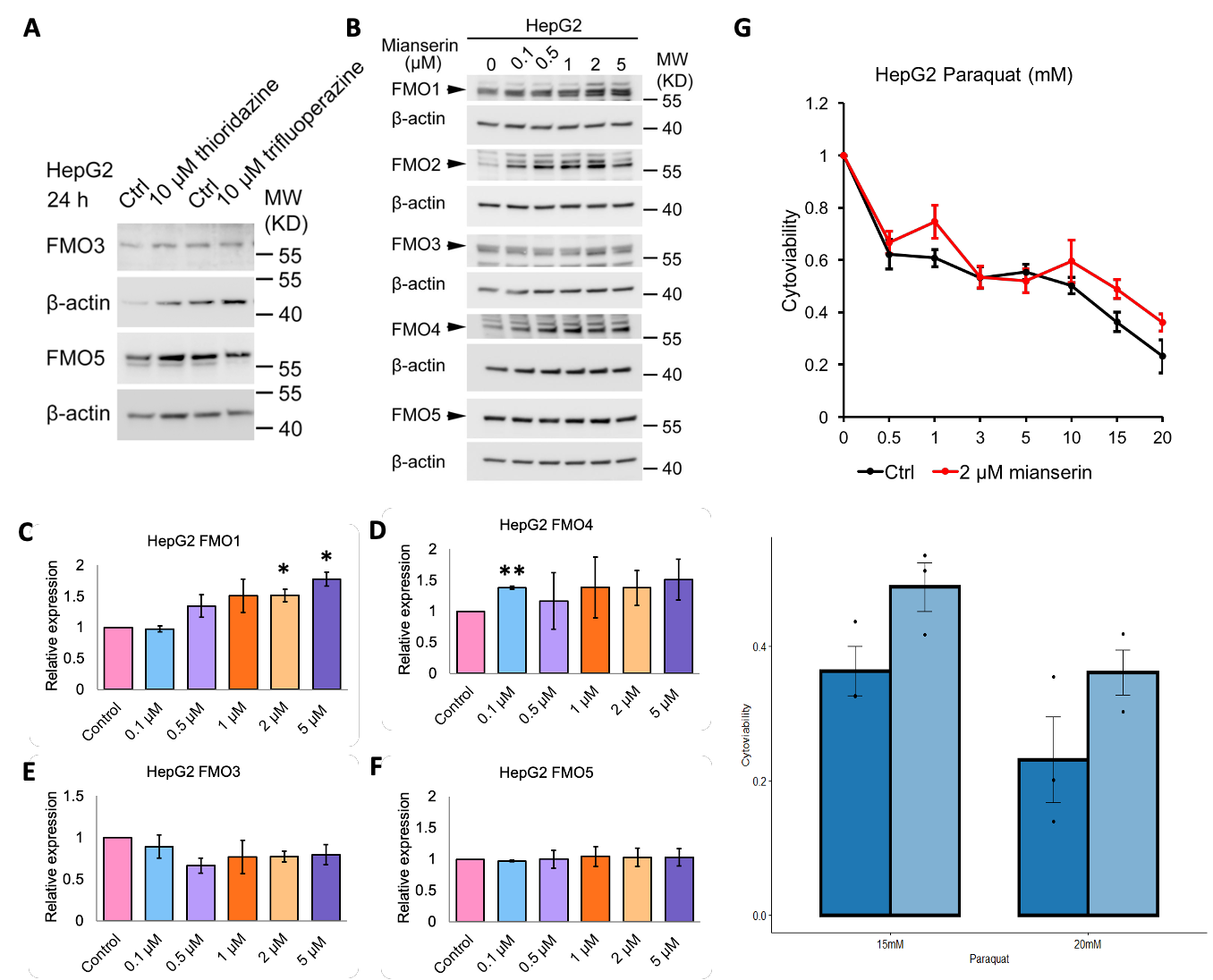

**Figure S9. Induction of Fmos by biogenic amine antagonists.** Representative western blot image (A) of FMO3 and FMO5 in whole cell lysates from HepG2 cells treated with 10 µM thioridazine or trifluoperazine. Representative western blot image (B) and quantification of FMO2 (C), FMO4 (D), FMO3 (E), and FMO5 (F) in whole cell lysates from HepG2 cells treated with 0.1 µM, 0.5 µM, 1 µM, 2 µM, or 5 µM mianserin. Cell survival percentages of HepG2 cells treated with 2 µM mianserin (light blue) or untreated (dark blue) under 15 mM or 20 mM paraquat (G). * denotes P<.05, ** denotes P<.01 when compared to control (Tukey’s HSD).

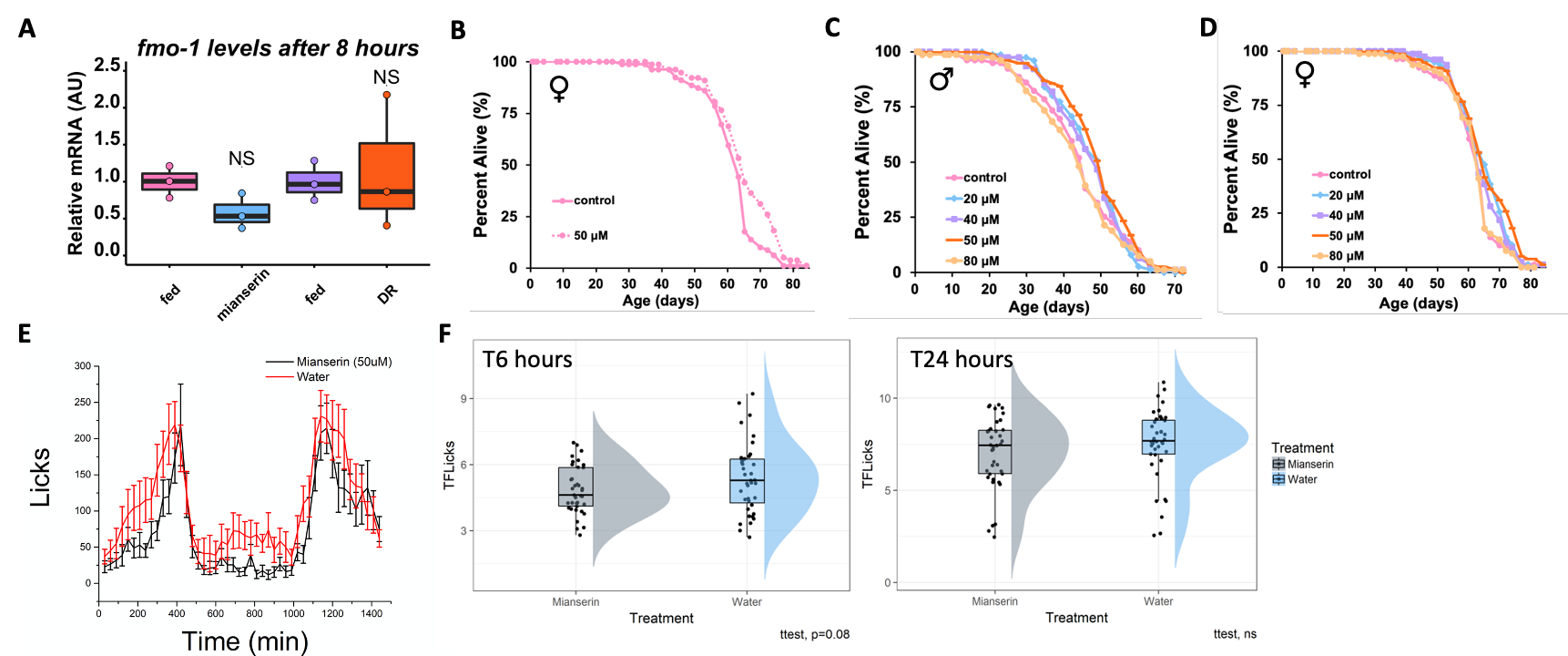

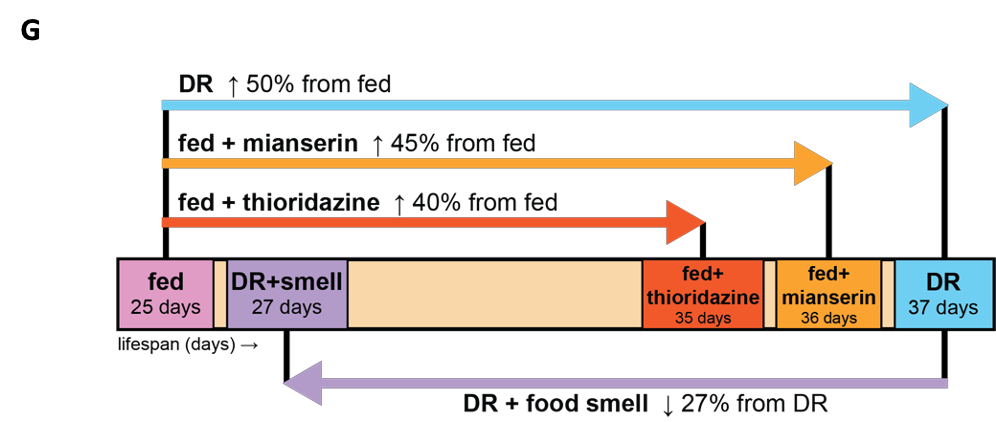

**Figure S10. Effects of DR mimetic mianserin on Fmo expression, feeding, and lifespan.** Fmo-1 mRNA levels (A) after 8 hours of 100µM mianserin (blue) or starvation (orange) compared to water controls (pink and purple, respectively) (red) or starvation (blue) compared to water control (black). Survival curves of female (B) flies dosed with water (solid line) or 50µM (dotted line) of mianserin. Combined survival curves of male (C) and female (D) flies dosed with water (pink), 20µM (blue), 30µM (purple), 40µM (orange), 50µM (yellow), or 80µM (purple) mianserin. 24 hours of FLIC assay data monitoring food intake in control (black) and mianserin (red) treatment (E). Extracted FLIC data (F) at 6 and 24 hours. Summary of the effects of food perception on DR and DR-mimetics longevity (G).

**References:**

1.	Fitch DHA. Introduction to nematode evolution and ecology. WormBook, ed. The C.

elegans Research Community: Wormbook; 2005.

2.	Frokjaer-Jensen C, Wayne Davis M, Hopkins CE, et al. Single-copy insertion of transgenes in Caenorhabditis elegans. 10.1038/ng.248. *Nat Genet*. 2008;40(11):1375-1383. doi:<http://www.nature.com/ng/journal/v40/n11/suppinfo/ng.248_S1.html>

3.	Berkowitz LA, Knight AL, Caldwell GA, Caldwell KA. Generation of stable transgenic C. elegans using microinjection. *J Vis Exp*. Aug 2008;(18)doi:10.3791/833

4.	Sutphin GL, Kaeberlein M. Measuring Caenorhabditis elegans life span on solid media. *J Vis Exp*. 2009;(27)doi:1152 [pii]

10.3791/1152

5.	Petrascheck M, Ye X, Buck LB. A high-throughput screen for chemicals that increase the lifespan of Caenorhabditis elegans. A*nn N Y Acad Sci.* Jul 2009;1170:698-701. doi:10.1111/j.1749-6632.2009.04377.x

6.	Zarse K, Ristow M. Antidepressants of the serotonin-antagonist type increase body fat and decrease lifespan of adult Caenorhabditis elegans. P*LoS One.* 2008;3(12):e4062. doi:10.1371/journal.pone.0004062

7.	Bargmann CI, Hartwieg E, Horvitz HR. Odorant-selective genes and neurons mediate olfaction in C. elegans. C*ell.* Aug 1993;74(3):515-27. doi:10.1016/0092-8674(93)80053-h

8.	Worthy SE, Haynes L, Chambers M, et al. Identification of attractive odorants released by preferred bacterial food found in the natural habitats of C. elegans. P*LoS One.* 2018;13(7):e0201158. doi:10.1371/journal.pone.0201158

9.	Linford NJ, Bilgir C, Ro J, Pletcher SD. Measurement of lifespan in Drosophila melanogaster. J *Vis Exp.* Jan 2013;(71)doi:10.3791/50068

10.	Murakami S, Salmon A, Miller RA. Multiplex stress resistance in cells from long-lived dwarf mice. F*ASEB J.* Aug 2003;17(11):1565-6. doi:10.1096/fj.02-1092fje
